# Transcriptional repression of *GTL1* under water-deficit stress promotes anthocyanin biosynthesis to enhance drought tolerance

**DOI:** 10.1101/2024.03.06.583754

**Authors:** Noel Anthony Mano, Mearaj A. Shaikh, Joshua R. Widhalm, Chan Yul Yoo, Michael V. Mickelbart

**Affiliations:** Department of Botany and Plant Pathology and Center for Plant Biology, Purdue University, West Lafayette, IN 47907, USA; Center for Plant Biology, Purdue University, West Lafayette, IN 47907, USA; Department of Horticulture and Landscape Architecture, Purdue University, West Lafayette, IN 47907, USA; School of Biological Sciences, The University of Utah, Salt Lake City, UT 84112, USA

**Keywords:** RNA-seq, transcription factor, polyamine, leaf development

## Abstract

The transcription factor GT2-LIKE 1 (GTL1) has been implicated in orchestrating a transcriptional network of diverse physiological, biochemical, and developmental processes. In response to water-limiting conditions, GTL1 is a negative regulator of stomatal development, but its potential role in other water-deficit responses is unknown. We hypothesized that GTL1 regulates transcriptome changes associated with drought tolerance over leaf developmental stages. To test the hypothesis, gene expression was profiled by RNA-seq analysis in emerging and expanding leaves of wild-type and a drought-tolerant *gtl1-4* knockout mutant under well-watered and water-deficit conditions. Our comparative analysis of genotype-treatment combinations within leaf developmental age identified 459 and 1073 differentially expressed genes in emerging and expanding leaves, respectively, as water-deficit responsive GTL1-regulated genes. Transcriptional profiling identified a potential role of GTL1 in two important pathways previously linked to drought tolerance: flavonoid and polyamine biosynthesis. In expanding leaves, negative regulation of *GTL1* under water-deficit conditions promotes biosynthesis of flavonoids and anthocyanins that may contribute to drought tolerance. Quantification of polyamines did not support a role for GTL1 in these drought-responsive pathways, but this is likely due to the complex nature of synthesis and turnover. Our global transcriptome analysis suggests that transcriptional repression of GTL1 by water deficit allows plants to activate diverse pathways that collectively contribute to drought tolerance.

## Introduction

Drought stress impacts plant growth and development via diverse physiological, biochemical, and developmental processes. Consequently, plants activate many distinct pathways to acclimate to water deficit. Water-deficit responsive transcription factors (TFs) are responsible for establishing a hierarchical transcriptional regulatory network for many of these responses, some being directly activated by tissue dehydration, others being downstream targets of signaling cascades controlled by other TFs. In plants, the DEHYDRATION RESPONSIVE ELEMENT BINDING (DREB) and APETALA2/ETHYLENE RESPONSIVE FACTOR (AP2/ERF) TFs are among the major and best characterized TF families in responding to abiotic stresses generally, and water-deficit conditions specifically (reviewed in Xie et al., 2019; Manna et al., 2021). Manipulation of TF expression results in a broad range of responses, with as few as 130 differentially expressed genes (DEGs) in a *NUCLEAR FACTOR Y* (*NFYA5*) overexpression line (Li et al., 2008) or as many as 9000 DEGs in a knockout mutant of *WRKY54* (Chen et al., 2017). For example, overexpressing members of the *NAC* TF family results in improved drought survival via longer roots, proline accumulation, and reduction in reactive oxygen species (ROS) accumulation, suggesting functional diversity among members of this TF family as well as the regulation of diverse pathways to facilitate drought tolerance (Figueroa et al., 2021).

One TF with a potentially broad range of regulatory targets is GT2-LIKE 1 (GTL1), a member of the plant-specific trihelix TF family (Zhou, 1999). GTL1 contains two trihelix DNA-binding domains (Breuer et al., 2009), through which GTL1 can modulate the expression of target genes. GTL1 represses the expression of genes that promote the transition between cell cycle phases (Breuer et al., 2009, 2012), cell extension (Shibata et al., 2018), and stomatal development (Yoo et al., 2010, 2019). As a result, *GTL1* knockout mutants have larger trichomes (Breuer et al., 2009), longer root hairs (Shibata et al., 2018), and fewer stomata (Yoo et al., 2010, 2019). Water-deficit stress activates Ca^2+^/calmodulin (CaM) that binds to GTL1 and allosterically inhibits DNA-binding activity, resulting in de-repression of *STOMATAL DENSITY AND DISTRIBUTION 1* (*SDD1*) (Yoo et al., 2019). The upregulation of *SDD1* and resulting inhibition of stomatal development results the *gtl1* knockout mutant having water-deficit tolerance via lower transpirational water loss (Yoo et al., 2010).

In Arabidopsis, GTL1 is a transcriptional regulator reported to have *ca*. 3900 putative direct DNA-binding targets and to act as a promoter of the light response and water and fluid transport but a repressor of the reactive oxygen species response (Breuer et al., 2012). Later, Shibata et al. (2018) showed that GTL1 and RSL4 formed a negative feedback loop in which GTL1 represses cellular signaling and cell wall biogenesis in roots. Finally, Völz et al. (2018) showed that GTL1 promoted bacterial immunity through the MAP kinase signaling and oxidative stress pathways. Transcriptomic experiments from trichomes, roots, and whole seedlings from nonfunctional *GTL1* mutants have been performed in well-watered conditions, but how they function under water-deficit conditions remains unexplored.

In Arabidopsis, the initial stages of leaf development involve almost completely proliferative growth, driven by the rapid division of leaf cells (Donelly et al., 1999; Andriankaja et al., 2012). This stage is characterized by the expression of the *GROWTH REGULATING FACTOR* (*GRF*) and *GRF INTERACTING FACTOR* (*GIF*) families (Horiguchi et al., 2005), cell cycling genes such as *CYCLIN DEPENDENT KINASE*s (Wang et al., 1997; Wang et al., 2000), and *OLIGOCELLULA 5/RIBOSOMAL PROTEIN 5* (*OLI5*/*RPL5A*) that regulate development of the ribosomal machinery (Fujikura et al., 2009). At this stage, the differentiation of leaf cell types such as guard cells, trichomes, and veins is initiated (Larkin et al., 1996; Geisler et al., 2000; Kang and Dengler, 2004; Andriankaja et al., 2012). Next, cell division ceases and cells transition to expansion growth, driven by cell wall loosening and turgor pressure, during which time the activity of ANGUSTIFOLIA and ROTUNDIFOLIA (Tsuge et al., 1996) balances length and width expansion. At the same time, jasmonic acid synthesis represses further mitotic division and induces phenylpropanoid metabolism (Pauwels et al., 2008), and genome replication occurs without cell division to increase cell size (endoreduplication) (Donelly et al., 1999; Beemster et al., 2005).

As leaf growth dynamics shift with time, certain specific drought tolerance processes have been identified to function differently in young and mature leaves, or are employed only at certain developmental stages. For example, young Arabidopsis leaves (about half the size of fully expanded leaves) accumulate more soluble sugars and proline, resulting in greater non-photochemical quenching and consequently more open photosystem II reaction sites, compared to fully expanded leaves under moderate drought conditions (Sperdouli and Moustakas, 2014). Under moderate drought, young leaves also accumulate more flavonoid compounds compared to fully expanded leaves (Sperdouli et al., 2021). Moreover, Skirycz et al. (2010) identified gene expression changes suggesting leaves composed of proliferating or expanding cells respond to mannitol-induced osmotic stress by upregulating biotic stress pathways: WRKY TFs, *MILDEW RESISTANT LOCUS O* (*MLO*) genes, and glucosinolate biosynthesis genes. Baerenfaller et al. (2012) described a reduction in carbon fixation genes in drought-stressed leaves across different growth stages, but did not explicitly contrast gene expression differences in drought-stressed leaves between the different stages.

One phenotype that is differentially sensitive to drought depending on leaf age is stomatal development. Since Arabidopsis stomatal development terminates fairly early in the life of a leaf (Geisler et al., 2000; Andriankaja et al., 2012), this pathway can only result in a different stomatal phenotype if drought imposition has occurred prior to stomatal development initiation in an examined leaf. When osmotic stress has been imposed in this manner, stomatal index (SI) in Arabidopsis decreases (Kumari et al., 2014; Yoo et al., 2019), indicating inhibition of stomatal development. This inhibition occurs through post-translational modification of the SPEECHLESS protein (Kumari et al., 2014) or via reduction of gene expression or calcium-calmodulin interaction with GT2-LIKE 1 (GTL1; Yoo et al., 2019). GTL1 promotes stomatal development via transrepression of *SDD1*, leading to a reduced stomatal phenotype, lower transpirational water loss, and thus higher survival under progressive drought compared to wild-type plants (Yoo et al., 2010). GTL1 also represses endoreduplication in trichome cells (Breuer et al., 2009), promotes salicylic acid metabolism leading to bacterial-triggered immunity (Völz et al., 2018), and represses root hair growth (Shibata et al., 2018). These diverse functions are possible because the GT3 box to which GTL1 binds is found in the promoter regions of many genes; therefore, in the absence of GTL1, many gene functional categories are differentially expressed (Breuer et al., 2012). Crucial to this study, among the categories of DEGs represented in the *gtl1* mutant was the response to abiotic stimuli.

Given the potential diversity of molecular processes regulated by GTL1, we propose that GTL1 regulates drought tolerance via molecular mechanisms distinct from stomatal anatomy. Given its role in the cell cycle and cell fate (stomatal development), we also hypothesized that GTL1 may differentially regulate gene expression in leaves that are in the cell division or cell expansion phases. We examined gene expression patterns in emerging and expanding leaves of wild-type and *gtl1-4* plants under well-watered and water-deficit conditions.

## Materials and Methods

### Plant growth, water-deficit imposition, and tissue collection

*Arabidopsis thaliana* plants of the Col-0 ecotype and the *gtl1-4* mutant were planted in a 1:2 soil mix of Turface (PROFILE Products LLC, Buffalo Grove IL) and Fafard F2 soilless media (Sungro Horticulture, Agawam MA) in SC7 Ray Leach ‘Cone-Tainers’ (Steuwe & Sons Inc, Tangent OR). Seeds were germinated and grown for the first 10 d after germination with overhead mist for 4 seconds every 6 minutes. When the first true leaves emerged, plants were transferred to a growth chamber (Conviron Inc, Winnipeg MB, Canada), and maintained under an 8-hour photoperiod, with a light intensity of 150 μmol m^-2^ s^-1^, day/night temperature and vapor pressure deficit of 23/18°C and 1.2/0.8 kPa, respectively, and relative humidity of 65%. After transferring plants to the growth chamber, they were photographed, and the length across the rosette was measured. Within both genotypes, rosettes beyond one quartile from the median size were removed from the experiment. The remaining seedlings were then randomized across the growth chamber in a completely randomized design. Plants were irrigated with acidified water supplemented with a water-soluble fertilizer (ICL Specialty Fertilizers, Dublin OH) to provide the following (in mg L^-1^): 150 N, 9.8 P, 119 K, 12 Mg, 21 S, 1.5 Fe, 0.4 Mn and Zn, 0.2 Cu and B, and 0.1 Mo. Nitrate and ammonium sources of nitrogen were provided as 61 and 39% total N, respectively. Irrigation water was supplemented with 93% sulfuric acid (Brenntag, Reading PA) at 0.08 mL L^-1^ to reduce alkalinity to 100 mg L^-1^ and pH to a range of 5.8 to 6.2.

All plants were irrigated and weighed to obtain the media-saturated weight (MSW). Subsequent irrigation was performed as needed to maintain well-watered plants between 65 and 100% media water content (MWC) and every two days to maintain water-stressed plants at 30% MWC (Supplemental Figure 1). MWC was calculated as [media fresh weight (MFW) - media dry weight (MDW)]/[MSW-MDW] × 100. During the experiment, a MDW of 49 g, based on dry weights obtained in prior experiments, was used to determine the appropriate amount of water to add to containers. At the conclusion of the experiment, containers were dried in a forced-air oven, and the actual MDW was obtained for each plant.

Plants were photographed every two days to identify leaf 14 that developed after the 30% target MWC was reached. Leaf area was quantified from images taken every 2 days. Leaf 14 was harvested from experimental plants 14 or 15 days after emergence in well-watered and water-deficit plants, respectively, when leaves were 50% of their estimated final size. At the same time, a day-old leaf (leaf 24–25) that had been identified in the previous day’s image was also harvested. Leaves were harvested into individual Eppendorf tubes and frozen in liquid nitrogen. In plants that were grown alongside RNA-seq plants but not harvested for RNA, leaves were tracked to full expansion, giving leaf area data from emergence to full expansion. Leaf area over time data was fitted with a sigmoidal curve (Figure 1a), which was then used to confirm that leaves harvested for RNA-seq were at the intended 50% of the estimated final leaf area.

**Figure 1.**
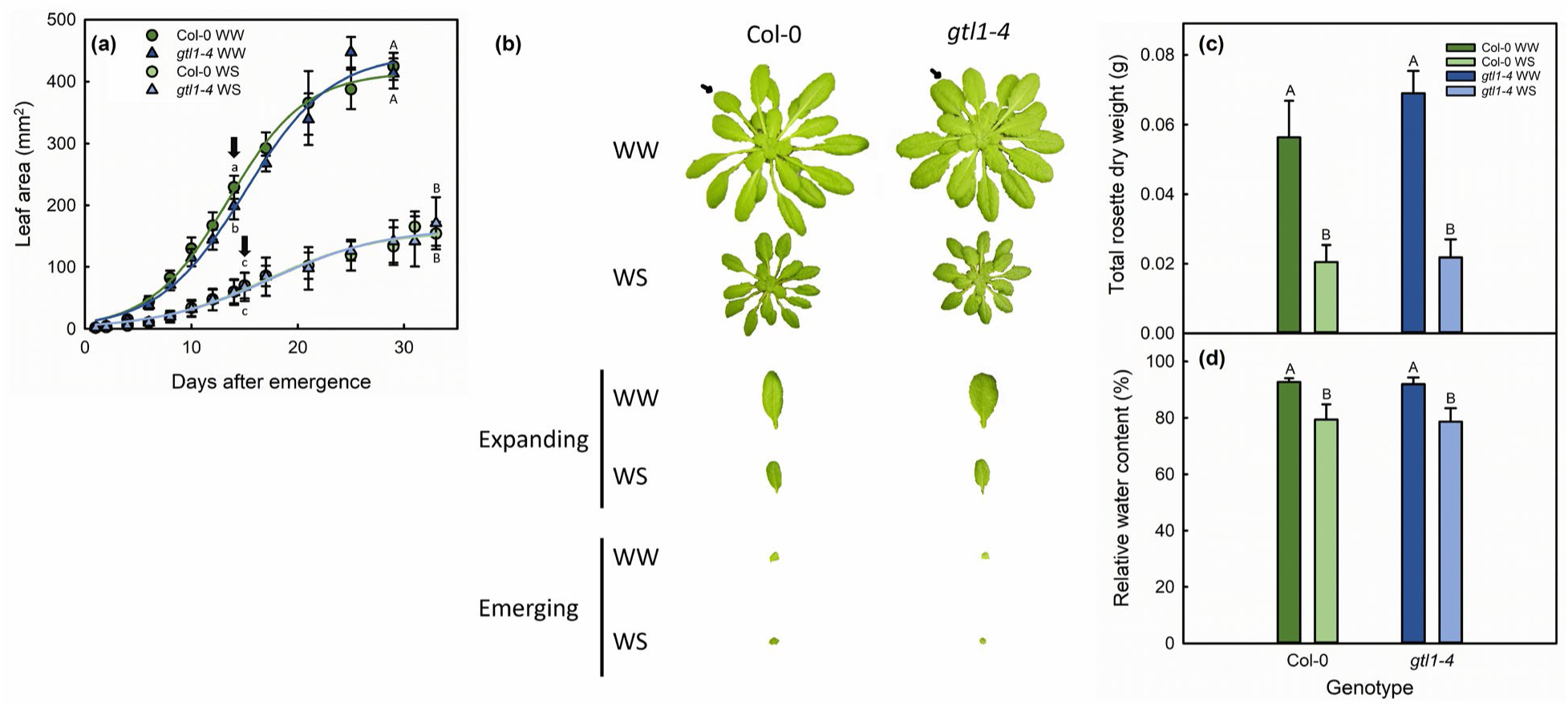
Leaf growth over time (a), representative whole rosettes and harvested expanding and emerging leaves (b), total rosette dry weight (c), and relative water content (d) of well-watered (WW, dark colored bars) and water-stressed (WS) in Col-0 and *gtl1-4* plants used for RNA-seq analysis. Different letters above bars or points represent statistically significant differences between genotype-treatment combinations at the *P* < 0.05 level. In (a), lowercase and uppercase letters are used for expanding and fully expanded leaves, respectively. Block arrows in (a) indicate leaf harvest times, and small arrows in (b) mark the expanding leaves collected from the rosettes. Bars represent standard error, *n =* 10 for rosette and RWC data, *n =* 4 for leaf size at full expansion.

Simultaneous to collecting leaf 14, leaf 13 was collected and weighed to obtain leaf fresh weight (LFW). Leaf 13 was subsequently placed in an 1.5-mL microcentrifuge tube with water and kept on a lab bench for 6 h, before being weighed to obtain the turgid weight (LTW). The leaf was then dried in an oven for 48 h before being weighed again to obtain the dry weight (LDW). Leaf relative water content (RWC) was calculated as [LFW-LDW]/[LTW-LDW] × 100. After leaves 13 and 14 were harvested, the whole rosette was cut at the root-shoot junction, segmented into individual leaves, and then bagged to be dried to weigh the total rosette dry weight.

After leaf 14 was fully expanded in the plants used to complete the leaf area curve, adaxial and abaxial surfaces were pressed onto cyanoacrylate droplets (Henkel Corporation, Düsseldorf, Germany) on glass slides to create impressions of leaf epidermis. Impressions were viewed under a BH-2 light microscope (Olympus, Center Valley PA) at 200X magnification, giving a field of view of 0.113 mm^2^. Four images were collected per field of view, and cell types were counted in ImageJ. Stomatal index was calculated as [number of stomata]/[number of stomata + number of epidermal cells].

### RNA extraction and sequencing

The experiment had three biological replicates with ten plants per genotype-treatment-development stage pooled per replicate. Total RNA was extracted from leaves using the QIAGEN PowerPlant RNA kit (QIAGEN, Germantown MD) following the manufacturer’s instructions. gDNA was removed from the RNA using the QIAGEN DNase Max Kit. RNA was then submitted to the Purdue Bioinformatics Core for quality assessment, library construction, and sequencing. A strand-specific paired-end read was performed with a read length of 50 bp and sequencing depth of 30 million reads/sample.

RNA used for RNA-seq analysis was also used to synthesize cDNA with the SuperScript III First-Strand Synthesis System (ThermoFisher Scientific, Waltham MA). qPCR was performed using the Luna Universal qPCR Master Mix (New England Biolabs, Ipswich MA) and the ThermoFisher StepOnePlus Real-Time PCR system to quantify the expression of *GTL1* and the leaf proliferation stage marker *ANGUSTIFOLIA 3* (*AN3*). Gene expression was normalized against that of the *ACTIN 2* (*ACT2*) housekeeping gene and quantified using the double-delta Ct method (Pfaffl, 2001). Primer sequences are given in Supplemental Table 1.

### Sequence data analysis

Sequence data were checked for quality with FastQC and low-quality reads (phred score ≤33) were removed with TrimGalore. Post trimming, the sequence files were then aligned to the *A. thaliana* genome (Araport 11 release) with STAR-aligner. The reference genome was prepared with the transcript quantification program RSEM, using the TAIR10 annotation of the genome. Diagnostics for the alignment step showed that the % of unaligned reads was ≤ 10% across all samples. The output bam files were then used to generate transcript counts per million (TPM) and expected counts using RSEM’s abundance estimation function. Expected count data was then passed to EBSeq for normalization and filtration of low-expression genes. Genes whose 75^th^ quantile of normalized expected counts were less than 10 were considered low-expression genes and were removed. This reduced the dataset from ∼37,000 genes to ∼24,000 and ∼23,000 genes in the emerging and expanding datasets, respectively. We used the EBSeq-HMM method (Leng et al., 2015) to generate a list of genes that were differentially expressed in response to water deficit and regulated by GTL1, using a Benjamini-Hochberg False-Discovery Rate of *q* < 0.05.

We filtered our DEG list as follows. First, genes that were differentially expressed in response to water deficit in Col-0 were identified. We next removed from consideration genes with similar directional fold changes to water deficit in Col-0 and *gtl1-4*, leaving 459 and 1073 genes in emerging and expanding leaves that were differentially regulated in response to water deficit and putatively regulated by GTL1. We identified GTL1-regulated genes that were potential targets of direct GTL1 binding by identifying GT3 boxes (5’-GGTAAA-3’) in the 2 kb region upstream of gene transcription start sites (TSS). GTL1-regulated genes were submitted to Ensembl’s Biomart database, querying the database for the region 0.5 and 2 kb upstream of the TSS. These sequences were downloaded and searched for the presence of a GT3 box.

### Secondary metabolite quantification

Secondary metabolites were quantified from expanding leaves of wild-type and *gtl1-4* plants grown under well-watered conditions similar to that of the RNA-seq experiment. Expanding leaves from 42-day-old plants were frozen and ground to a fine powder in liquid nitrogen. The following protocol was derived from procedures previously used to quantify total flavonoids and anthocyanins in leaf tissues (Zhang et al., 2018; Wade et al., 2003; Yu et al., 2019; Zhang et al., 2019; Havaux and Kloppstech, 2001). Ground powder (*ca*. 40 mg FW) was added to 2 mL 1% HCl in methanol and left at 4°C in darkness for 24 hours. Chloroform (2 mL) and deionized water (1 mL) were added, and the mixture was mixed vigorously. The upper phase was taken for measurement of absorption at 530 and 600 nm using a Beckman Coulter DU730 UV-Vis spectrophotometer (Beckman Coulter Life Sciences, Indianapolis IN). The following equation was used to calculate the concentration of total anthocyanins as follows (Yu et al., 2019):

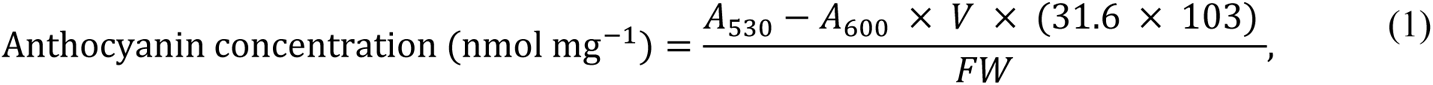

where *A*530 and *A600* are absorbances at 530 and 600 nm, respectively, *V* is the extraction volume, (31.6 x 103) is the extinction coefficient, and *FW* represents the fresh weight of the powdered leaf sample (mg). Flavonoid concentration was determined using a standard curve of 1 mg mL^-1^ quercetin in 1% HCl-methanol, and therefore, is expressed as ng quercetin equivalent per mg powdered leaf fresh weight.

The polyamines putrescine, spermine, and spermidine were quantified from an average of 69 mg of the powdered tissue left over from quantifying total flavonoids and anthocyanins. Polyamines were quantified using a protocol adapted from Anwar et al. (2019) and Herrero et al. (2016). We used a Zorbax SB-C18 5 μm 4.6 x 250 mm column at 35°C on a 1260 Infinity HPLC instrument with a fluorescence detector (Agilent Technologies, Santa Clara CA) using an excitation wavelength of 340 nm and emission wavelength of 510 nm. Standard curves were generated using analytical standards of putrescine, spermine, spermidine, and the internal standard 1,7-diaminoheptane (Sigma-Aldrich, Inc., St. Louis MO).

## Results & discussion

### Water stress resulted in reduced leaf expansion and stomatal development

The sustained water-deficit conditions in this experiment resulted in water-stressed plants experiencing a consistent MWC of 30% (Supplemental Figure 1). Well-watered plants were watered to media saturation as MWC approached 70%, preventing over-watering while maintaining an average MWC of 85%. There were no differences in MWC between genotypes within a given treatment, so wild-type Col-0 and *gtl1-4* experienced identical water supply regimes (Supplemental Figure 1). Growth data further supported that the two genotypes were identically water stressed; Col-0 and *gtl1-4* leaves were of similar size when harvested for RNA-seq and at full expansion (Figure 1a). *gtl1-4* rosettes had shorter petioles and leaves had larger trichomes compared to wild-type plants, but otherwise, the two genotypes were visually similar (Figure 1b). Col-0 and *gtl1-4* plants grown under water-deficit stress were smaller (Figure 1b, c), and leaves had lower relative water content (Figure 1d). Water content and rosette growth reductions occurred to a similar degree in both genotypes. Together, the data indicate that *gtl11-4* and Col-0 growth responded identically to sustained water deficit, which ensured that leaves of both genotypes were at the same growth stage and stress level for transcriptome profiling. Therefore, observed differences between the two genotypes can be attributed to differential genetic responses to water deficit without being confounded by growth stage or stress severity differences. Consistent with previous findings (Yoo et al., 2019), GTL1 promotes stomatal development and *gtl1-4* plants lacked the reduction of stomatal index observed in water-stressed Col-0 plants (Supplemental Figure 2).

We sampled 1-day-old (emerging) and 50% of final leaf area (expanding) leaves (Figure 1a, b) to contrast these roles of GTL1. Expected count data was normalized and filtered to remove non- or low-expressed genes across all genotype-treatment combinations. This procedure was performed separately within emerging and expanding leaves and resulted in 24311 and 22858 genes that could be quantified in each leaf type. Within emerging and expanding leaves, the EBSeq-HMM procedure (Leng et al., 2015) was used to identify DEGs across all genotype-treatment combinations. This procedure enabled direct comparison of gene expression levels, avoiding the risk of false positives that would have arisen from comparing lists of DEGs from multiple one-to-one contrasts.

We first assessed expected differences in specific gene expression across the different tissue stages and water-deficit levels. We selected 12 common drought-stress markers, including abscisic acid (ABA)-dependent and ABA-independent signaling. Nine of these twelve genes were upregulated as expected (Supplemental Figure 3). Many of these drought-stress markers were initially identified under conditions of progressively withheld water (Baker et al., 1994) or leaf dehydration assays (Yamaguchi-Shinozaki and Shinozaki, 1993; Liu et al., 1998), and the lack of differential expression we observed in some drought-stress markers is consistent with previous observations using a sustained water deficit (Harb et al., 2010; Baerenfaller et al., 2012; Rymaszewski et al., 2017). Based on the consistent upregulation of water-deficit markers, smaller leaf size, and lower leaf relative water content (Figure 1), we can conclude that plants of both genotypes were water-deficit stressed.

We also examined the expression of fifteen genes upregulated in emerging leaves (Stage 1 from Baerenfaller et al. (2012)), and fifteen genes upregulated later in leaf development (Stage 2 and in the transition from Stage 2 to 3). Thirteen of fifteen markers of the emerging leaf stage were upregulated in emerging leaves relative to expanding leaves (Supplemental Figure 4). In our dataset, all fifteen markers of the expanding leaf stage were upregulated in expanding leaves relative to emerging leaves. There were no genotype differences in the expression of these marker genes. Additionally, qPCR- and RNA-seq-based quantification of the leaf proliferation stage marker *AN3* (Horiguchi et al., 2005) showed much higher expression in emerging leaves than in expanding leaves (Supplemental Figure 5). Therefore, our dataset had a clear distinction between leaves in the emerging stage compared to older leaves that were undergoing expansion growth.

### Water deficit downregulates GTL1 expression in emerging and expanding leaves

In emerging leaves, *GTL1* expression was downregulated in response to water deficit (Figure 2). RNA-seq detected log2 fold change of -0.5 of *GTL1* expression in response to water deficit, whereas qPCR did not show a significant change in expression. A stronger degree of *GTL1* downregulation was observed in expanding leaves in response to water deficit. RNA-seq detected -0.71-fold *GTL1* downregulation due to water deficit, and qPCR also identified lower *GTL1* expression in water-stressed expanding leaves relative to well-watered controls. In both emerging and expanding *gtl1-4* leaves, RNA-seq detected some residual *GTL1* transcript sequence, but this was 3–4-fold lower compared to Col-0 plants. The four genotype-treatment combinations thus established a range of *GTL1* expression, with water-stressed wild-type plants as an intermediate between well-watered Col-0 and the *gtl1-4* plants, consistent with results reported previously by Yoo et al. (2010).

**Figure 2.**
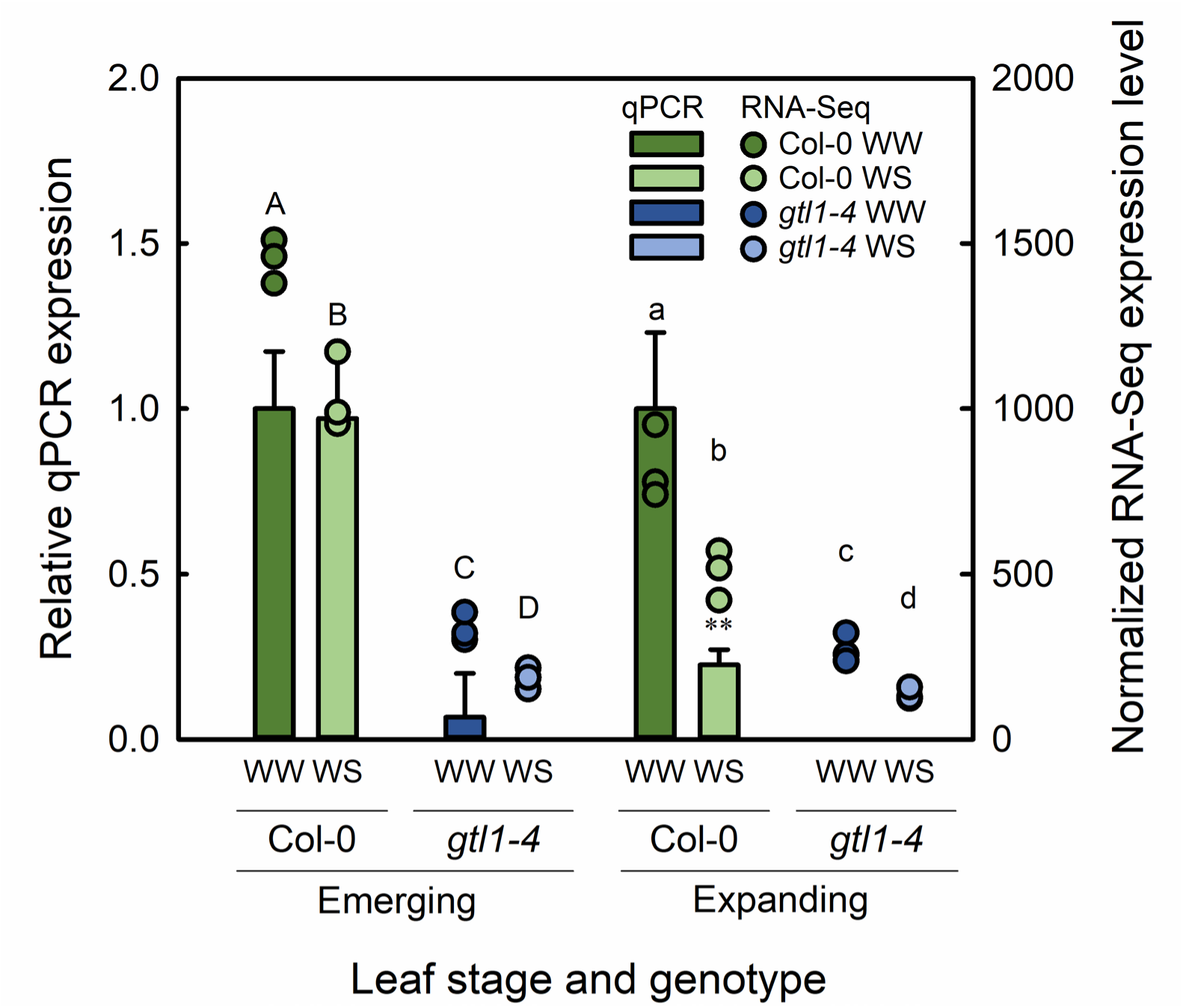
Expression of *GTL1* as quantified by qPCR (bars ± standard error, left axis) or RNA-seq (dots, right axis) in well-watered (WW) or water-stressed (WS) leaves. Data were compared within leaf types. Different letters above bars indicate mean separation as tested with EBSeq-HMM. qPCR data was compared against well-watered Col-0, and ** indicates a statistically significant difference at *P* < 0.01 (*n =* 3).

### GTL1-regulated water-deficit genes were identified

EBSeq-HMM allowed us to directly compare genotype-treatment combinations within leaf developmental stages and thus identify genes that were differentially expressed in response to water deficit in Col-0 and regulated by GTL1. Genes that had the same expression level between the two genotypes in well-watered and water-stressed conditions were not considered to be GTL1-regulated. After filtering for non-or-low-expressed genes, 24311 and 22858 genes could be analyzed in emerging and expanding leaves (Table 1). In emerging leaves, 1962 genes were upregulated in response to water deficit and 2360 genes were downregulated. Most of these genes were similarly expressed in both genotypes under well-watered and water-stressed conditions. Therefore, they were not considered to be GTL1-regulated. In emerging leaves, 154 and 305 genes up- and-downregulated in response to water deficit were GTL1-regulated (totaling 459 GTL1-regulated genes, shown in the overlapping white area of Figure 3a). In expanding leaves, 1503 genes were upregulated, and 1704 genes were downregulated in response to water deficit. Of these, 530 upregulated genes and 543 downregulated genes were GTL1-regulated (totaling 1073 GTL1-regulated genes, shown in the overlapping white area of Figure 3b). Therefore, from the different genotype-treatment combinations, our dataset resulted in 459 and 1073 GTL1-regulated water-deficit responsive genes in emerging and expanding leaves (white regions in Figure 3a, b). The fact that *GTL1* expression was more strongly downregulated in expanding leaves than in emerging leaves (Figure 2) explains the higher number of GTL1-regulated water-deficit responsive genes in expanding leaves compared to emerging leaves.

**Figure 3.**
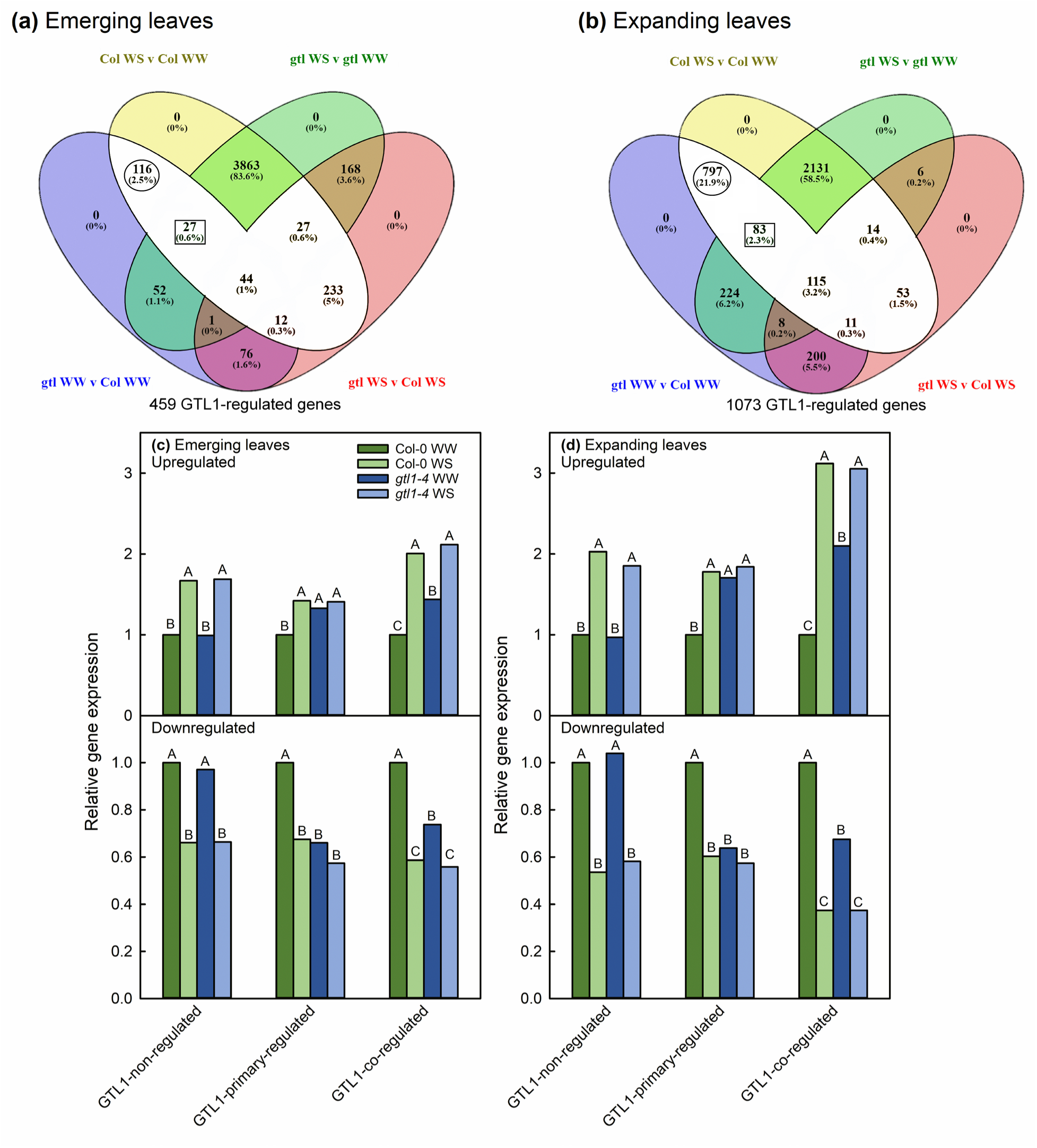
Venn diagrams of differentially expressed genes in specific genotype-treatment combinations for emerging (a) and expanding leaves (b). The region filled in white are responsive to water deficit in wild-type plants but regulated by GTL1 (bolded in Table 1 and indicated as totals beneath each Venn diagram). GTL1-regulated genes were primary-regulated by GTL1 (circled) or co-regulated by GTL1 and water deficit (outlined in a square). Gene regulation patterns in response to water deficit and GTL1 are shown for emerging (c) and expanding leaves (d), based on average relative gene expression to well-watered wild-type leaves. Mean separation values are as identified by EBSeq-HMM.

**Table 1.** Number of genes that were or were not differentially regulated in response to water stress (WS) as well as GTL1-regulated in the different leaf development stages.

| Leaf type | Genes differentially expressed in response to WS in wild-type plants |  |  |  |
| --- | --- | --- | --- | --- |
|  | Upregulated |  | Downregulated |  |
|  | Not GTL1-regulated | GTL1-regulated | Not GTL1-regulated | GTL1-regulated |
| Emerging | 1808 | **154** | 2055 | **305** |
| Expanding | 972 | **530** | 1159 | **543** |

Bolded numbers are genes which are differentially expressed under WS and also GTL-1 regulated. These genes are the focus of further analyses.

We identified a subset of GTL1-regulated water-deficit responsive genes as responsive to water deficit via promotion or repression by GTL1 (Figure 3c, d). These genes (circled in Figure 3a, b) had an expression level in well-watered *gtl1-4* that was identical to that of water-stressed Col-0. Therefore, for these genes, the absence of GTL1 effectively phenocopied their expression in water-stressed conditions (Supplemental Figure 6a, d). We refer to these as GTL1-primary-regulated genes. These genes were the majority of GTL1-regulated water-deficit responsive genes in expanding leaves (797 out of 1073), but only a quarter of GTL1-regulated water-deficit responsive genes in emerging leaves (116/459). We also observed that genes could be additively promoted or repressed by GTL1 and water deficit, and we refer to these as GTL1-co-regulated genes (Figure 3c, d).

First, we performed GO analysis exclusively on GTL1-primary-regulated genes. However, this analysis found relatively little evidence of functional category overrepresentation (Supplemental Figure 6). Next, we performed GO analysis on all GTL1-regulated water-deficit responsive genes, which broadened the representation of GO categories. To include genes that were both completely and additively regulated by GTL1 and water deficit, subsequent pathway analysis was performed using the GO results from the pooled GTL1-primary-regulated and GTL1-co-regulated genes. Expanding leaves will be the focus of the remainder of our analysis because GTL1-regulated genes in over-represented GO categories could be mapped to previously characterized molecular pathways.

### GTL1 promotes protein synthesis via ribosome biogenesis and organization

In emerging leaves, endoplasmic reticulum (ER) body organization was over-represented among GTL1-regulated genes upregulated in response to water deficit (Figure 4a). The group of ER body organization genes consisted of *PYK10* and *NAI2*, the latter of which was previously identified as a direct binding target of GTL1 (Breuer et al., 2012). The gene product of *PYK10* is a *β*-glucosidase involved in hydrolyzing indole glucosinolates. In Arabidopsis, ER bodies contain large amounts of *β*-glucosidases, among which PYK10 is the most abundant (Nakano et al., 2017). PYK10 is functionally dependent on *NAI2* expression because NAI2 is required for ER body formation (Yamada et al., 2008).NAI2 was also identified as a negative regulator of growth and proline accumulation in drought-stressed plants (Kumar et al., 2015). In our emerging leaf dataset, *NAI2* and *PYK10* expression were upregulated under water-deficit conditions only in Col-0. Therefore, the repressive effect of NAI2 on the drought response would have been mitigated in *gtl1-4*. *PYK10* may be upregulated in water-stressed plants as indole glucosinolates are a key component of pathogen resistance and defense (reviewed in Burow and Halkier (2017)). Therefore, the lack of *PYK10* differential expression may explain the compromised immunity of *gtl1* plants to *Pseudomonas syringae* observed by Völz et al. (2018). However, since there is a weak correlation between the transcript and protein level of *PYK10* (Yamada et al., 2008), a GTL1-mediated PYK10-NAI2 mechanism that represses water deficit in Col-0 plants is highly speculative and needs future molecular and biochemical characterization.

**Figure 4.**
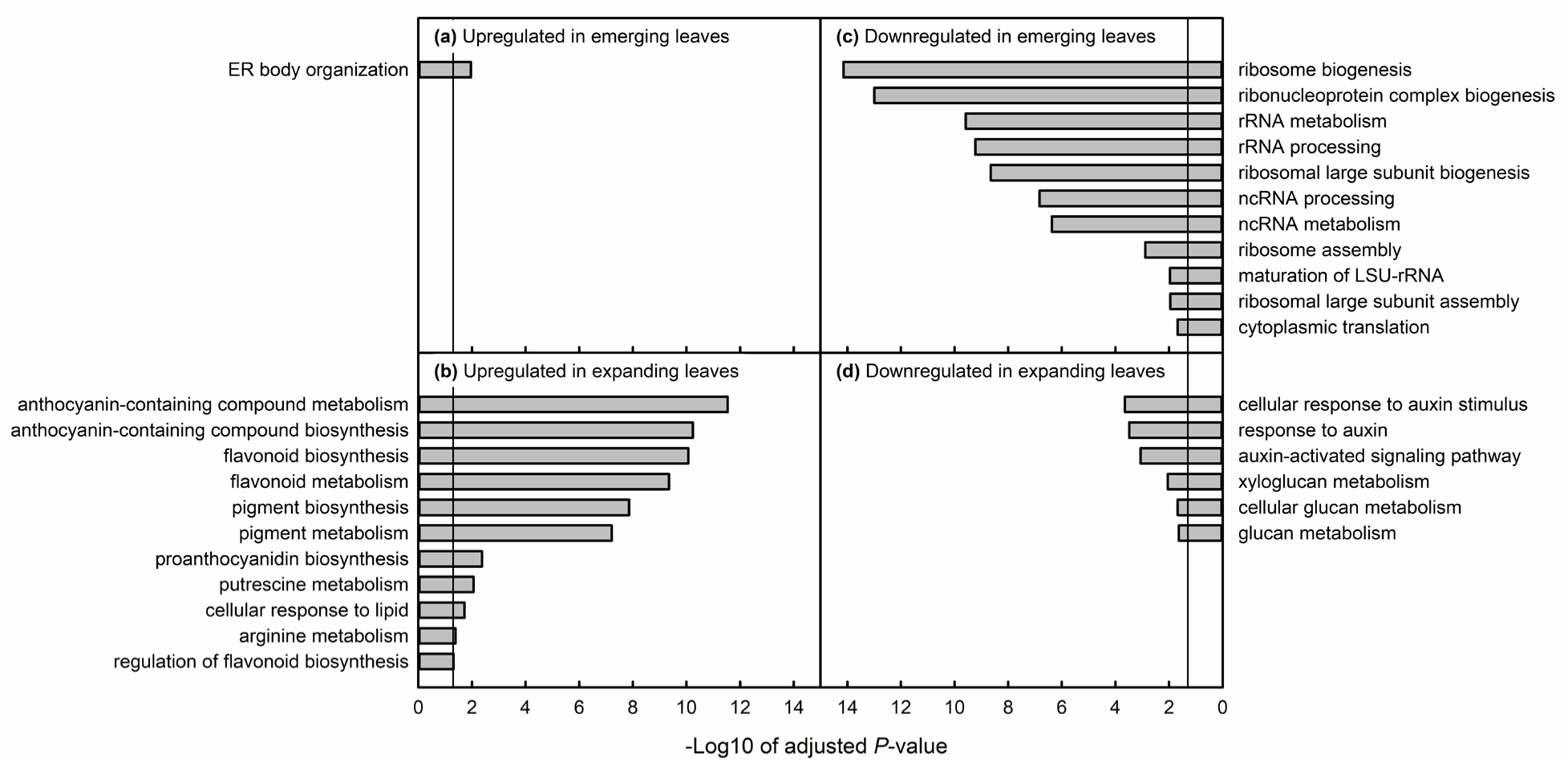
Results of gene ontology overrepresentation analysis for genes that were upregulated (a, b) or downregulated (c, d) in response to water deficit and regulated by GTL1. GO analysis was performed for all GTL1-regulated water-deficit responsive genes (white regions in Figure 3). All gene categories for which overrepresentation was statistically significant are shown. The vertical line in each plot represents the threshold of the false-discovery rate statistical significance.

In emerging leaves, ribosome biogenesis and organization were over-represented functional categories among GTL1-regulated genes downregulated in response to water deficit (Figure 4c, Supplemental Figure 7). Many of these genes were not characterized beyond their functional annotation, so it was not possible to analyze the contribution of specific GTL1-regulated ribosomal genes to drought tolerance. However, modifying ribosome biogenesis has knock-on effects on plant development (Rosado et al., 2010; Wieckowski and Schiefelbein, 2012) and potential stress tolerance. Sormani et al. (2011) demonstrated that transcriptional regulation of nuclear ribosomal protein genes is correlated with the eventual localization of the ribosomal protein product to the cytoplasm, mitochondria, or plastid. Alongside the ‘ribosome code’ proposed by Komili et al. (2007), the differential expression of ribosome biogenesis genes creates a mechanism by which the level of translation in the cytoplasm is decreased in response to stress. Differential expression (Martinez-Seidel et al., 2020) or sequestration of ribosome component transcripts (Merret et al., 2017) results in the production of different ribosome complexes in temperature-stressed plants. Accessions of Arabidopsis have qualitatively diverse pools of ribosomal rRNA (Rabanal et al., 2017), although the functional significance of ribosomal diversity is not known. If many ribosomal biogenesis genes are downregulated prior to water-deficit stress in *gtl1-4*, new leaves in *gtl1-4* could, by way of differential protein expression, be in a drought-tolerant state even under well-watered conditions. Similarly, since proteins are energetically expensive to produce (Baerenfaller et al., 2012), reduced protein synthesis in well-watered *gtl1-4* could redirect carbon flux toward producing metabolites through drought-responsive pathways. Young leaves are more translationally active than older leaves (Baerenfaller et al., 2012; Omidbakhshfard et al., 2021), explaining why ribosome biogenesis/organization genes were over-represented in emerging but not expanding leaves.

### GTL1 represses pigment and polyamine biosynthesis and promotes the auxin response pathway

In expanding leaves, genes involved the flavonoid and anthocyanin biosynthesis and the putrescine pathway were over-represented among GTL1-regulated genes upregulated under water deficit (Figure 4b). We were able to map the DEGs in each GO category onto each pathway (Figures 5, 6) since both are well-characterized biochemical pathways. TRANSPARENT TESTA 4 (TT4) catalyzes the conversion of coumaroyl-CoA to naringenin chalcone, the first committed step of the flavonoid and anthocyanin pathway (Spribille and Forkmann, 1982). *TT4* was transcriptionally repressed by GTL1, such that in both WW and WS *gtl1-4,* its expression was similar to that of water-stressed Col-0 (Figure 5a). *TT4* was the first of several GTL1-regulated flavonoid or anthocyanin biosynthesis genes to have a GT3 box within 2 kb upstream of its transcriptional start site (TSS), potentially indicating direct regulation of its expression by GTL1. Genes that encoded proteins for almost all the subsequent steps of flavonoid and anthocyanin biosynthesis were additively repressed by GTL1 or water-sufficient conditions, so their expression was partially upregulated in WW *gtl1-4*, then further upregulated to Col-0 WS levels in water-stressed *gtl1-4*. The TTG1-EGL3/TT8-PAP1 protein complex promotes the expression of downstream anthocyanin and flavonoid biosynthesis genes (Xu et al., 2014). Genes regulating the formation of this complex or encoding individual components of the complex were also repressed by GTL1 and then upregulated under water deficit. Supporting the genetic data, we found that concentrations of total flavonoids and anthocyanins were higher in well-watered expanding leaves of *gtl1-4* than those of Col-0 (Figures 5b, c). Expression of *TT9* was also downregulated and *GSTF12* was upregulated in *gtl1-4* and in response to water deficit. The *TT9* gene product traffics flavonoids to the vacuole (Ichino et al., 2014), whereas the same function is performed for anthocyanins by GSTF12 (Sun et al., 2012). Thus, GTL1 may not only repress the synthesis of flavonoids and anthocyanins but also maintain the vacuolar balance of these compounds.

**Figure 5.**
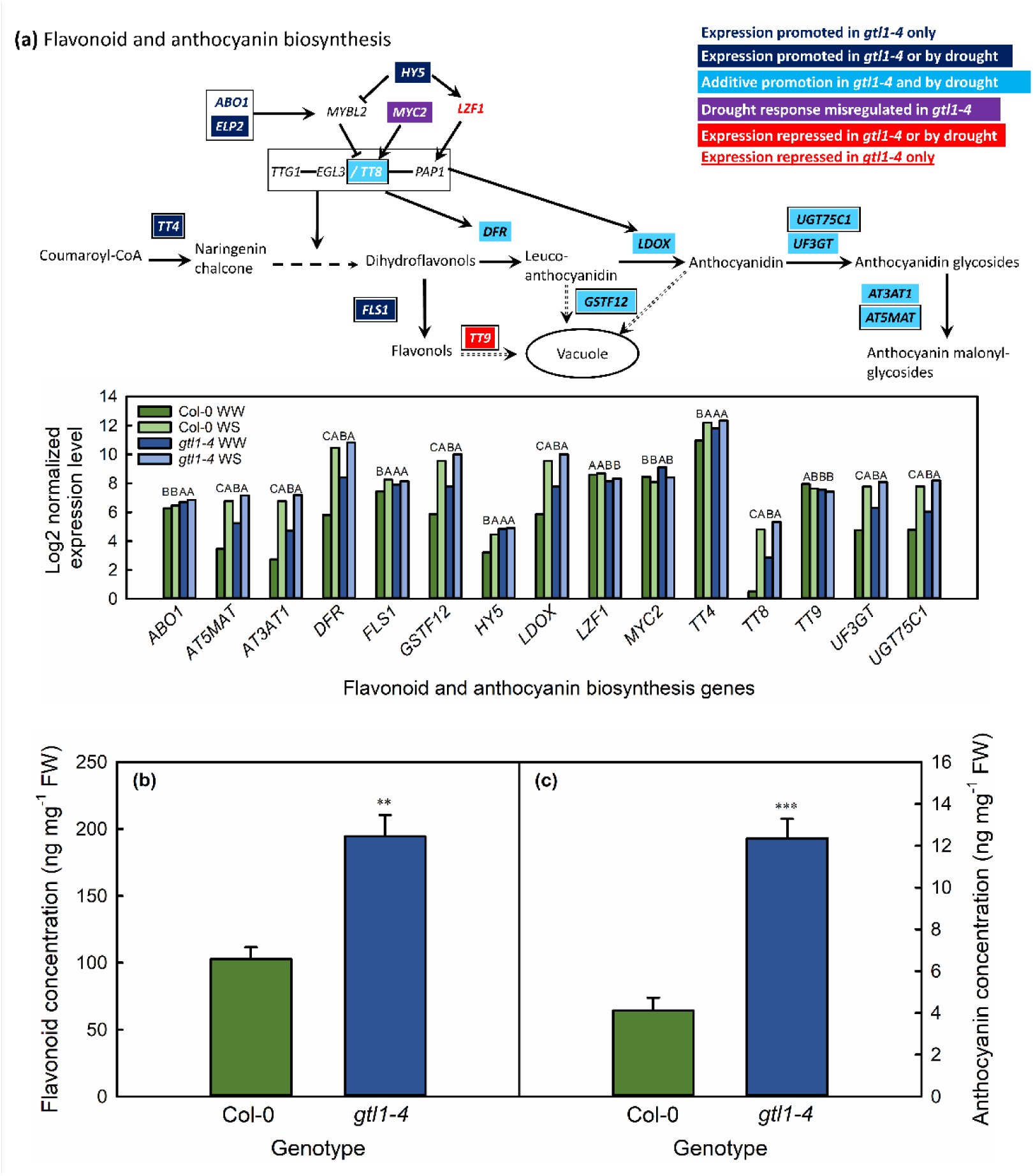
GTL1 regulated genes in the flavonoid and anthocyanin biosynthesis pathway (a), total flavonoids expressed as ng quercetin equivalent (b) and total anthocyanins (c) in well-watered expanding leaves of Col-0 and *gtl1-4*. Genes are colored according to their responsiveness to GTL1 and water deficit. Multiple boxed genes together encode protein products that form complexes. Genes with a GT3 box within 2 kb upstream of the transcriptional start site are indicated with a black border. Expression level of identified genes are shown as the log2-transformed normalized expression levels for three biological replicates per genotype in well-watered (WW) and water-stressed (WS) leaves. Statistically significant differences between genotype-treatment combinations are as given for *q* < 0.05 using the EBSeq-HMM package. For the biochemical data, ** and *** indicate statistically significant differences between genotypes as tested with a one-way ANOVA and post-hoc Tukey test.

Breuer et al. (2012) quantified downregulation by GTL1 of several flavonoid and anthocyanin biosynthesis genes (*FLS3*, *PAP3*, *PAL4*, *MLO4*, and *UGT71C4*) as being downregulated by GTL1. However, possibly due to the analysis being conducted in the trichomes of well-watered plants, these functional categories as a whole were not overrepresented (Breuer et al., 2012). Flavonoids and anthocyanins are both well-characterized in their ability to alleviate oxidative damage to plant cells via scavenging reactive oxygen species (ROS) (Nagata et al., 2003; Tohge et al., 2005; Nakabayashi et al., 2014; Xu et al., 2017; Wang et al., 2016). Arabidopsis ecotypes with high concentrations of anthocyanins accumulate lower concentrations of H_2_O_2_ under mild water-deficit conditions (Chen et al., 2021). ROS scavenging by flavonoids and anthocyanins promotes survival in drought and osmotically stressed plants, possibly because plants with increased flavonoid accumulation minimized water loss relative to wild-type plants (Nakabayashi et al., 2014). Xu et al. (2017) proposed that the anthocyanin accumulation response to stress is dependent on the later stages of the anthocyanin biosynthetic pathway, from the synthesis of leucocyanidin onwards, which was also reflected in drought-stressed ICE63 (Chen et al., 2021), and supported by the mechanism that has emerged in *gtl1-4*. Our dataset thus showed that under water-sufficient conditions, GTL1 represses several flavonoid and anthocyanin genes that play important roles in drought tolerance. The *gtl1* knockout mutant thus accumulates drought-protective metabolites during water-sufficient periods that may at least partially explain the observed drought tolerance of this mutant.

We also found that GTL1 represses polyamine biosynthesis genes in expanding leaves (Figure 6). Most of these DEGs were repressed by water-sufficient conditions via GTL1, such that the expression was similarly upregulated in WW and WS *gtl1-4* and WS Col-0 plants relative to WW Col-0. The exceptions to this expression pattern were *ADC2*, which was additively repressed by water sufficiency and GTL1, and *PAO4*, which was promoted by GTL1. The difference in GTL1 regulation of *PAO4* and *PAO2* may reflect the substrate specificities of the two enzymes: PAO4 metabolizes spermine, whereas PAO2 metabolizes both spermine and spermidine (Takahashi et al., 2010). *PAO4* and *PAO2* both had GT3 boxes 2 kb upstream of the TSS and are direct targets of GTL1 binding (Breuer et al., 2012). The observed additive repression of *ADC2* would facilitate the accumulation of agmatine (Watson et al., 1998), a putrescine precursor. The expression of genes for the reverse reactions of polyamine metabolism, that is, the synthesis of spermidine from putrescine (*SPERMIDINE SYNTHASE* (*SPDS*) *1*,*2*, or 3) or the synthesis of spermine from spermidine (*SPERMINE SYNTHASE*) were either not differentially regulated between the two genotypes or did not change in response to water deficit. The expression of *PAO1*, which encodes an enzyme that synthesizes 4-amino-butanal from spermidine (Takahashi et al., 2010), was similarly unresponsive to water deficit or genotype. Overall, the transcriptional data suggested the accumulation of putrescine at the expense of spermine and spermidine pools in *gtl1-4* and water-stressed Col-0.

**Figure 6.**
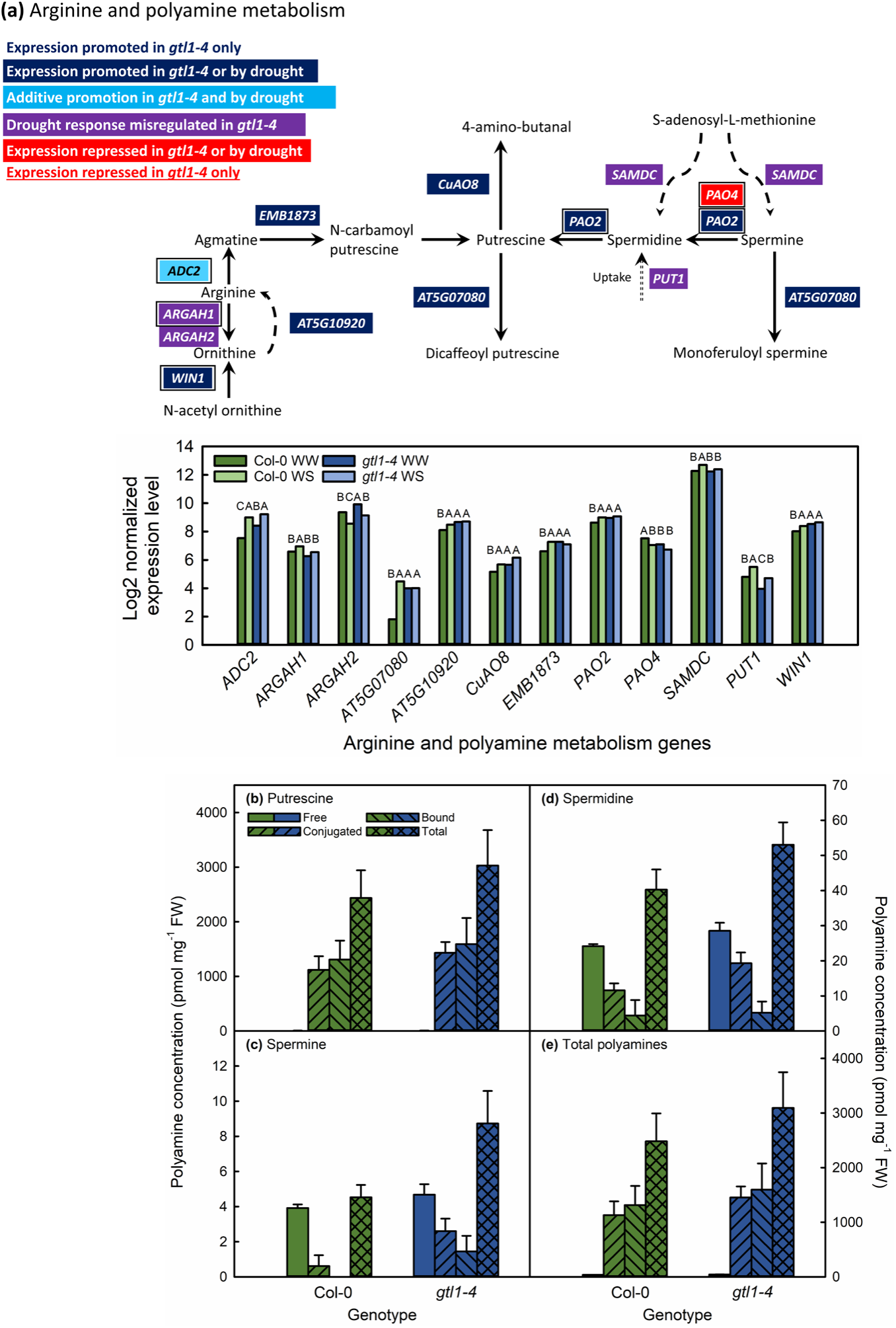
GTL1-regulated genes in the arginine and polyamine metabolism pathway (a), concentrations of free, conjugated, bound, and total putrescine (b), spermine (c), spermidine (d) and total polyamines (e) in well-watered expanding leaves of Col-0 and *gtl1-4*. Genes are colored according to their responsiveness to GTL1 and water deficit. Multiple boxed genes together encode protein products that form complexes. Genes with a GT3 box within 2 kb upstream of the transcriptional start site are indicated with a black border. Expression level of identified genes are shown as the log2-transformed normalized expression levels for three biological replicates per genotype in well-watered (WW) and water-stressed (WS) leaves. Statistically significant differences between genotype-treatment combinations are as given for *q* < 0.05 using the EBSeq-HMM package. No statistically significant differences were observed in the amount of free, conjugated, or bound polyamines between Col-0 and *gtl1-4*.

Of these three major polyamines in plants, putrescine may facilitate drought tolerance by promoting the scavenging of ROS (Wu et al., 2016). As we observed, salt and drought stress resulted in the upregulation of *ADC* and, subsequently the accumulation of putrescine (Urano et al., 2004; Alcázar et al., 2006). Therefore, transgenic plants overexpressing *ADC2* produced more putrescine and had a higher survival rate under drought stress relative to wild-type plants (Alcázar et al., 2010), with the opposite being true in plants with inhibited *ADC2* expression (Wu et al., 2016). Our transcriptional data indicated that spermine and spermidine would not accumulate during drought or in the absence of *GTL1* expression. This is supported by previous data indicating that except for *SPDS1*, spermine and spermidine synthesis genes either did not respond to water deficit or decreased to the expression level seen in unstressed plants (Alcázar et al., 2011).

In quantifying putrescine, spermine, and spermidine, we did not observe a difference in the amount of these individual or total compounds between Col-0 and *gtl1-4*, nor were there differences in the relative allocation of individual polyamines between free, bound, or conjugated forms (Figures 6b–e). Therefore, our biochemical data did not support the hypothesized increase in putrescine concentrations in *gtl1-4*. There are several possible explanations for this observation. First, *S-ADENOSYLMETHIONINE DECARBOXYLASE* (*SAMDC*), which encodes the enzyme that synthesizes the precursor to spermine and spermidine, was slightly upregulated in well-watered *gtl1-4* relative to well-watered Col-0 (Figure 6a). *SAMDC* is also post-transcriptionally regulated (Hanfrey et al., 2001; Franceschetti et al., 2001) based on polyamine levels (Ivanov et al., 2010). Therefore, *SAMDC* expression or protein levels may have helped maintain the consistency of spermine and spermidine concentrations between the two genotypes. Second, the upregulation of *PAO* and *EM1873*, responsible for the biosynthesis of putrescine and spermidine, was minimal (< 0.5 log2-fold change) compared to the >2 log-2 upregulation of *AT5G07080* (Figure 6a), which encodes the enzyme that converts putrescine and spermine to their dicaffeoyl and monoferuloyl derivatives (Wang et al., 2021). It is possible that the unchanged polyamine pool between *gtl1-4* and Col-0 is due to the production of phenolamides derived from polyamines since we did not measure flux through this metabolic pathway. There is limited data on the potential contributions of phenolamides towards water-deficit tolerance specifically (Roumani et al., 2021), but these compounds may play a role in ROS scavenging (Ding et al., 2019). If validated alongside elevated flavonoid and anthocyanin content of *gtl1-4*, there would thus be dual biochemical mechanisms to reduce oxidative damage in water-stressed plants of this mutant.

### GTL1 represses auxin-related genes

In expanding leaves, genes in the auxin response pathway were over-represented among GTL1-regulated genes that were downregulated in response to water deficit (Figure 4d). GTL1 promoted many genes throughout the auxin pathway, including biosynthesis, transport, and auxin-regulated phenotypes. Auxin biosynthesis inhibitor *SPL2*, as well as the biosynthesis gene *YUC8* were promoted by GTL1 and water-sufficiency, so the expression of these genes is similar between *gtl1-4* and water-stressed Col-0 (Figure 7). *GH3.17*, which conjugates auxin into an inactive auxin-amino conjugate (Staswick et al., 2005), is also promoted by GTL1 in the same way. These imply that in well-watered *gtl1-4* plants, active auxin levels may be reduced, potentially explaining the shortened petioles of *gtl1-4* (Figure 1). Auxin results in the remobilization of PIN proteins, which are polar transporters of auxin. GTL1 and water sufficiency repressed the expression of the chaperone *P23*, which facilitates PIN protein remobilization (D’Alessandro et al., 2015), and *TOR* (Yuan et al., 2020), which stabilizes PIN proteins. At the same time, expression of *PIN1* and *NDL3*, which encodes an auxin efflux protein (Mudgil et al., 2009), were downregulated to Col-0 WS levels in *gtl1-4*. Therefore, with our transcriptional data, we propose that GTL1 promotes auxin biosynthesis, homeostasis, and sensitivity under water-sufficient conditions. Our data supports previous investigations into the role of GTL1, first in the identification of auxin-responsive genes among GTL1 transcriptional targets in trichomes (Breuer et al., 2012), and auxin-regulated root hair development (Shibata et al., 2018). Additionally, Li et al. (2023) showed the auxin response pathway GO category as upregulated when the yellowhorn *GTL1* was overexpressed in well-watered conditions.

**Figure 7.**
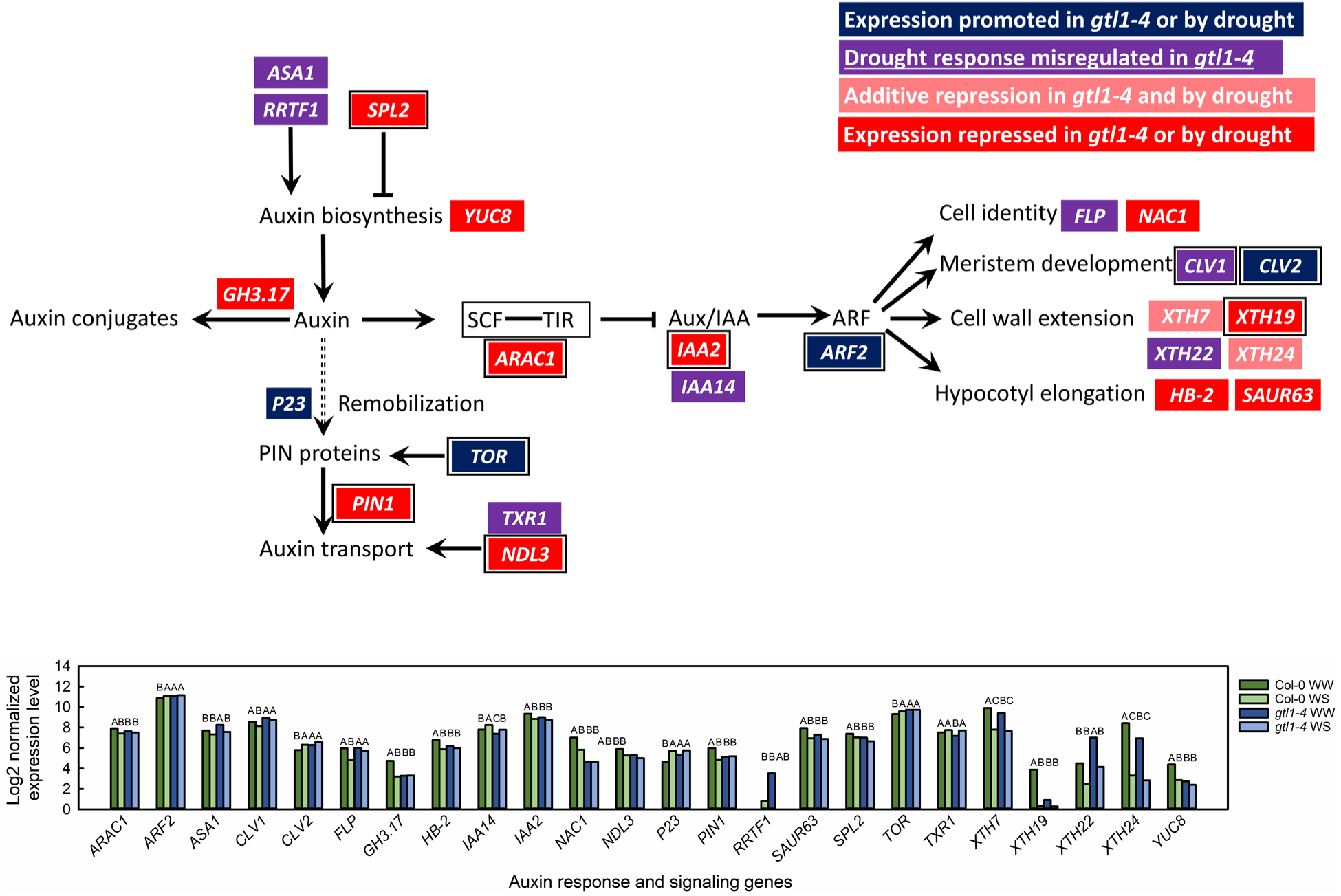
GTL1-regulated genes in the auxin response pathway. Genes are colored according to their responsiveness to GTL1 and water deficit. Multiple boxed genes together encode protein products that form complexes. Genes with GT3 boxes within 2 kb upstream of the transcription start site are indicated with a black border. Expression level of identified genes are shown as the log2-transformed normalized expression levels for three biological replicates per in well-watered (WW) and water-stressed (WS) leaves. Statistically significant differences between genotype-treatment combinations are as given for *q* < 0.05 using the EBSeq-HMM package.

In the absence of GTL1, the auxin response pathway may be inhibited. *ARAC1*, which recruits Aux/IAA proteins for degradation (Tao et al., 2005), and *IAA2* were both downregulated in *gtl1-4* and in water-stressed Col-0. Since Aux/IAA proteins repress AUXIN RESPONSE FACTORs, upregulation of *ARF2* in *gtl1-4* and water-stressed Col-0 was consistent with *IAA* downregulation. However, many downstream processes of auxin signaling were downregulated in *gtl1-4* or in response to water deficit. These especially included genes that promote cell wall extension and hypocotyl elongation (Figure 7). Cell expansion is limited under water-deficit and auxin levels (Shi et al., 2014) and signaling are often inhibited by water deficit (Kodaira et al., 2011; Chen et al., 2012). Ultimately, the role of GTL1 on the inhibition of auxin signaling by water deficit will need to be evaluated with *in vivo* characterization of auxin sensitivity under control and water-deficit conditions.

### Additional roles of GTL1 in regulating water-deficit response via analysis of individual genes

Next, we considered individual candidates to promote drought tolerance among the 116 and 797 GTL1-primary-regulated genes from emerging and expanding leaves, respectively. *TIP2;2* encodes an aquaporin, which, when not expressed, results in plants with higher drought survival (Feng et al., 2018). *TIP2;2* expression was reduced in *gtl1-4* and in response to water deficit. *SFP1* encodes a sugar transporter (Quirino et al., 2001) with a known downregulation response to water deficit (Slawinski et al., 2021). As with *TIP2;2*, we observed downregulation of *SFP1* in *gtl1-4* and water-stressed plants. *MYO-INOSITOL-1-PHOSPHATE SYNTHASE 2* (*MIPS2*) was repressed by GTL1 and water deficit and so was upregulated in WW and WS *gtl1-4* and in WS wild-type plants. MIPS2 catalyzes the synthesis of *myo-*inositol (Donahue et al., 2010), which accumulates under water-deficit conditions (Georgii et al., 2017; Li et al., 2020). The introduction of maize *MIPS* enhanced *myo*-inositol accumulation and drought tolerance in Arabidopsis (Li et al., 2020), possibly because of improved water relations in cells accumulating *myo*-inositol. *CYCLIN H;1* is also a well-characterized negative regulator of drought tolerance. Lower *CYCH;1* expression resulted in lower stomatal aperture, a lower rate of water loss, and higher ROS production (Zhou et al., 2013). While our emerging leaves were too young for the stomatal mechanism, ROS production can act as an abiotic stress signal (Miller et al., 2009). Therefore, downregulation of *CYCH;1* in young *gtl1-4* leaves may result in upregulated stress signaling under well-watered conditions.

In expanding leaves, many GTL1-primary-regulated genes had already been characterized earlier as part of the flavonoid/anthocyanin, putrescine, or auxin signaling pathways. Nevertheless, we additionally identified several genes that may facilitate water-deficit tolerance in *gtl1-4*. The TF *MYB102* is upregulated in response to osmotic stress, but its expression is directly repressed by AGL16. The knockout mutant *myb102* had lower seed germination and root growth under osmotic stress conditions (Zhao et al., 2021). *ARABIDOPSIS HALOTOLERANCE 2-LIKE* (*AHL*) encodes a 3′-phosphoadenosine-5′-phosphate (PAP) phosphatase. A T-DNA knockout of *AHL* results in a lower accumulation of the osmoprotectant proline, which in turn inhibits plant growth under osmotic stress (Shin et al., 2019). Overexpression of the rice orthologue *OsAHL1* improved rice survival and yield under osmotic and salt stress (Zhou et al., 2016). A more direct role in drought tolerance is found in *ABSCISIC ALDEHYDE OXIDASE 3* (*AAO3*), which encodes the enzyme that catalyzes the final step in ABA biosynthesis (Seo et al., 2000, 2004). *AAO3* expression is induced in response to water deficit (Koiwai et al., 2004; Endo et al., 2008) as part of ABA biosynthesis induction, and *aao3* mutants had lower survival under water-deficit conditions (Khan et al., 2019). All three of *MYB102*, *AHL*, and *AAO3* were transcriptionally repressed by GTL1, and so their constitutive upregulation in *gtl1-4* may facilitate a mild water-deficit response even under well-watered conditions. Putative *AAO3* and ABA accumulation *gtl1-4* explains the observed expression pattern of *HOMEOBOX PROTEIN 6* (*HB6*), which, as a GTL1-primary-promoted gene, was downregulated in response to water deficit and in WW and WS *gtl1-4*. The HB6 protein is a negative regulator of ABA sensitivity and ABA-induced stomatal closure (Himmelbach et al., 2002; Lechner et al., 2011). Therefore, downregulation of *HB6* may promote ABA-mediated tolerance of water deficit, particularly if ABA synthesis is validated to be increased in *gtl1-4*.

### Conclusions

We show here that GTL1 represses genes in flavonoid, anthocyanin, and polyamine synthesis. The gene expression data in the flavonoid and anthocyanin biosynthetic pathways were supported by increased accumulation of total flavonoids and total anthocyanins in expanding leaves of *gtl1-4* relative to that of wild-type plants. These pathways have been repeatedly characterized as facilitating tolerance to water-deficit conditions, so *GTL1* knockout results in a partially activated drought response in well-watered plants. Differential expression of most genes in these pathways was not the result of direct GTL1 binding, so GTL1 regulation is mediated by control of upstream TFs that are not yet identified. Many GTL1-regulated water-deficit responsive genes in emerging leaves have not yet been functionally characterized. However, GTL1 represses genes that facilitate ribosome biogenesis and organization, which may lead to a different proteome in young *gtl1-4* leaves. Our results thus suggest several mechanisms that explain drought tolerance in *gtl1-4* beyond the previously observed stomatal phenotype. Future work should test and attempt to experimentally validate these proposed drought tolerance mechanisms. Since GTL1 is functionally conserved in wheat (Zheng et al., 2016), poplar (Weng et al., 2012; Liu et al., 2021), and yellowhorn (Li et al., 2023), this study identifies drought tolerance mechanisms that may be a target for crop or silvicultural improvement.

## Acknowledgements

The authors wish to thank Mike Gosney, Nathan Deppe, and Dan Little for assistance with plant care and growth chamber maintenance. We are also grateful to Valerio Cirillo for assistance with data collection, and to Paul Hasegawa, Dan Szymanski, Yun Zhou, Clint Chapple, and Chao Cai for suggestions and advice on RNA-seq data analysis. Finally, we are grateful to Phillip San Miguel and Allison Sorg of the Purdue Bioinformatics Core for assistance and advice on RNA-seq sample preparation and sequencing. This project was funded by a BARD Grant to M.V.M., by the USDA National Institute of Food and Agri-culture Hatch Projects 1013618 to M.V.M. and 177845 to J.R.W., and by support from the Purdue Center for Plant Biology to N.A.M.

## Author contributions

N.A.M and M.V.M. designed the experiments. N.A.M. collected and analyzed the data. N.A.M., C.Y.Y., and M.V.M. wrote the paper. M.S. and J.R.W. designed and analyzed the polyamine experiment. M.S. and N.A.M. performed polyamine measurements. All authors read and edited the paper.

---

**Supplemental Table S1.**
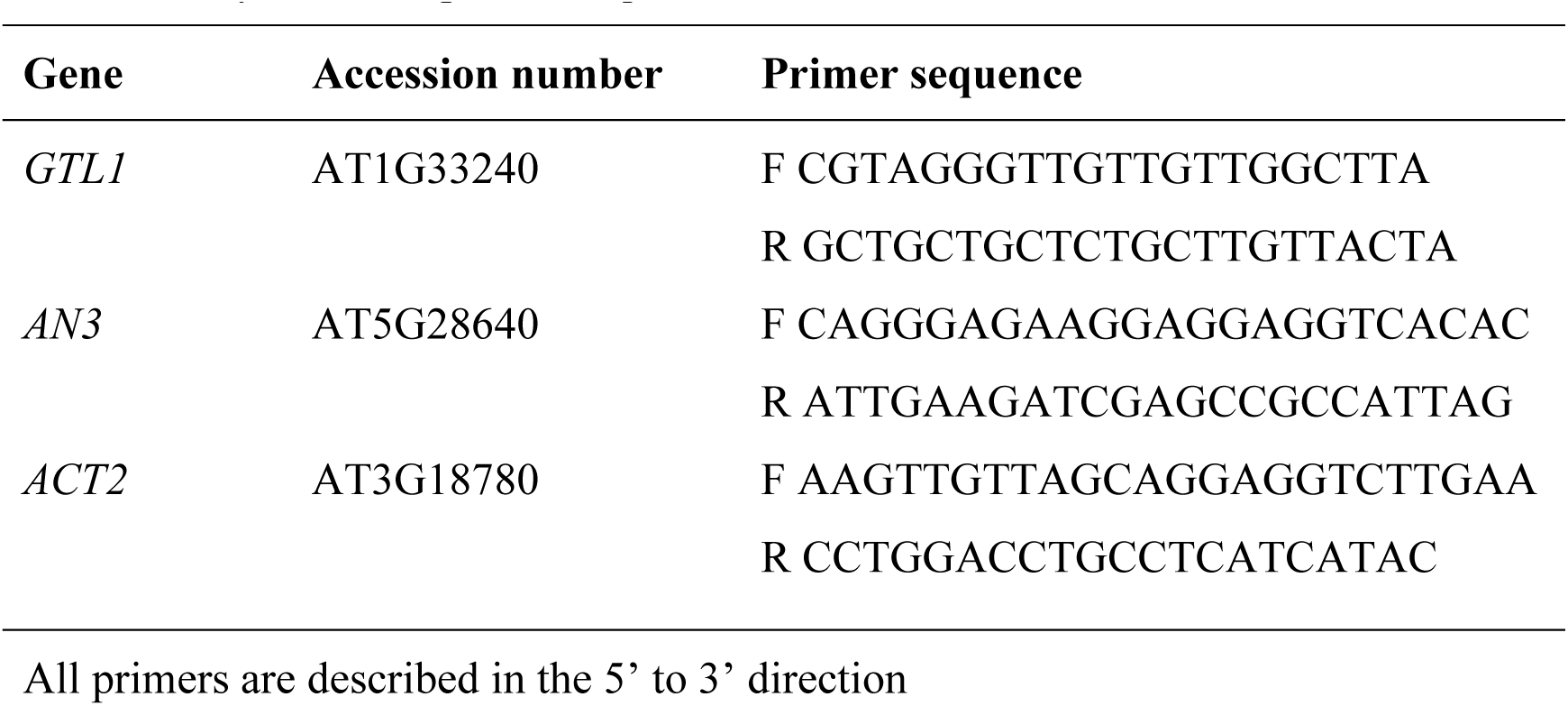
Accession numbers of genes quantified by qPCR in the current study and their primer sequences.

**Supplemental Figure 1.**
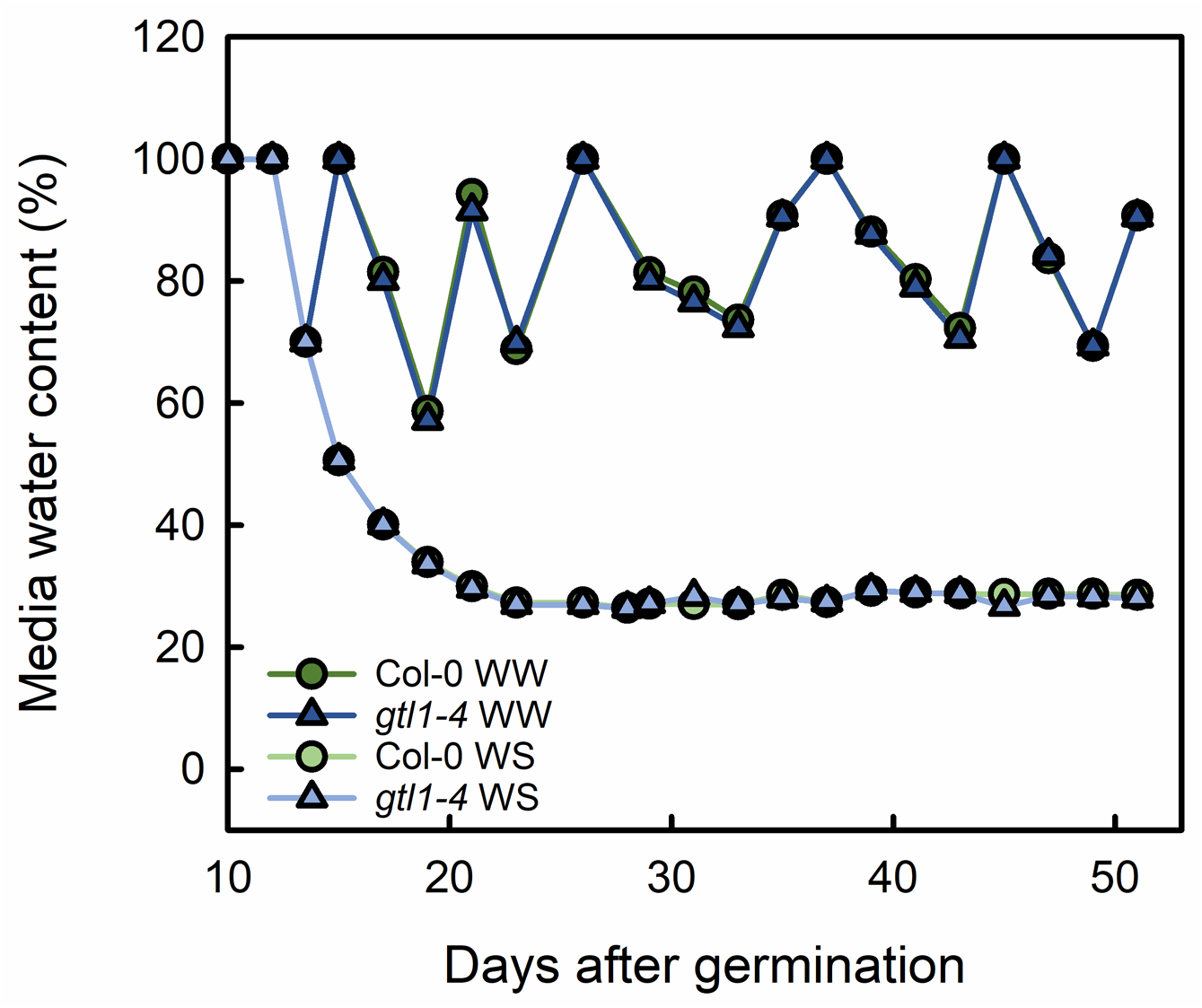
Average media water content over each irrigation interval of Col-0 and *gtl1-4* plants for the duration of the experiment.

**Supplemental Figure 2.**
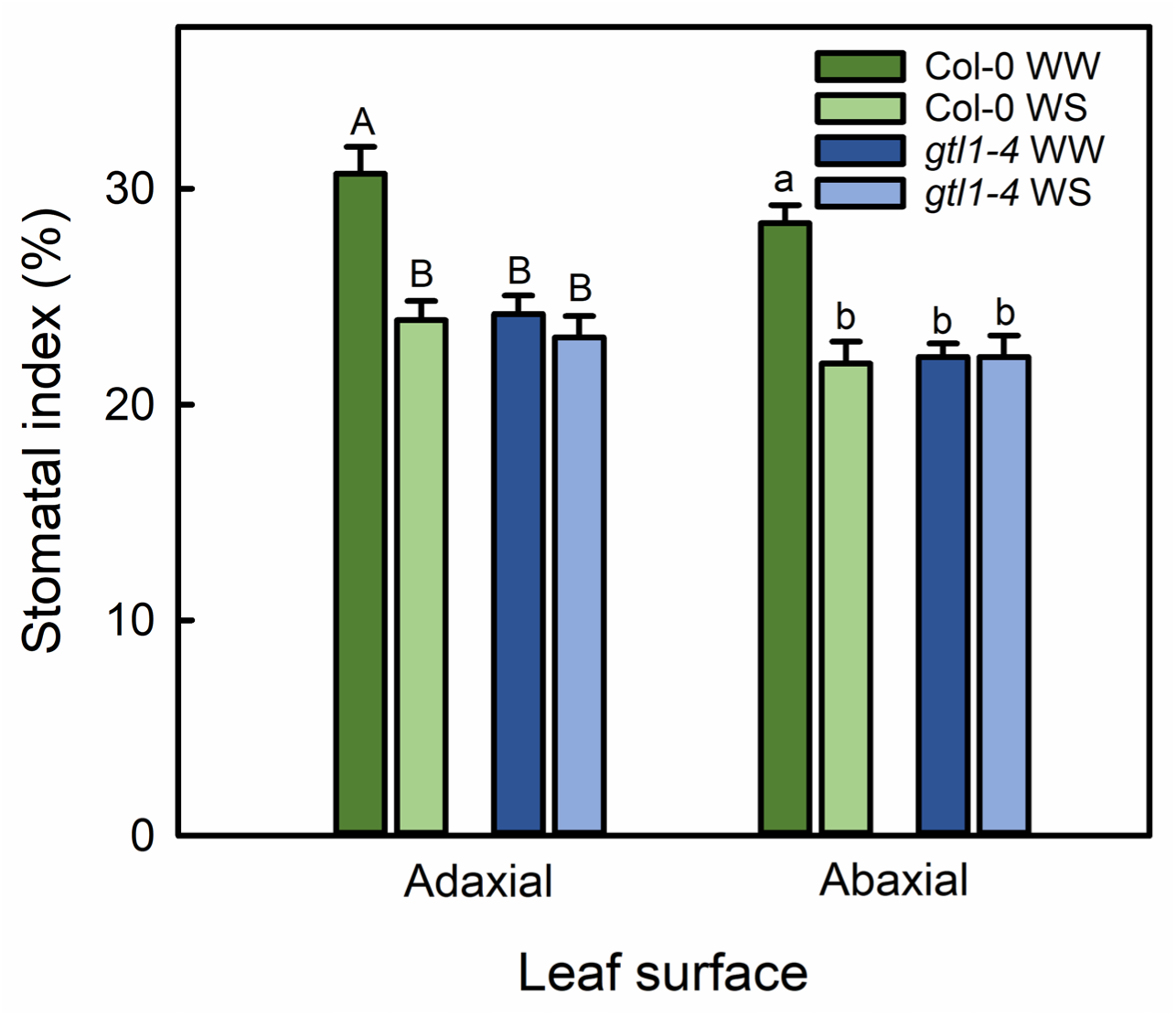
Stomatal index on the adaxial and abaxial surface in fully expanded well-watered (WW) and water-stressed (WS) Col-0 and *gtl1-4* leaves. Different letters above the bars show statistically significant differences at *P*<0.05 between genotypes and treatment groups within each leaf surface. Error bars represent the standard error, *n=* 8.

**Supplemental Figure 3.**
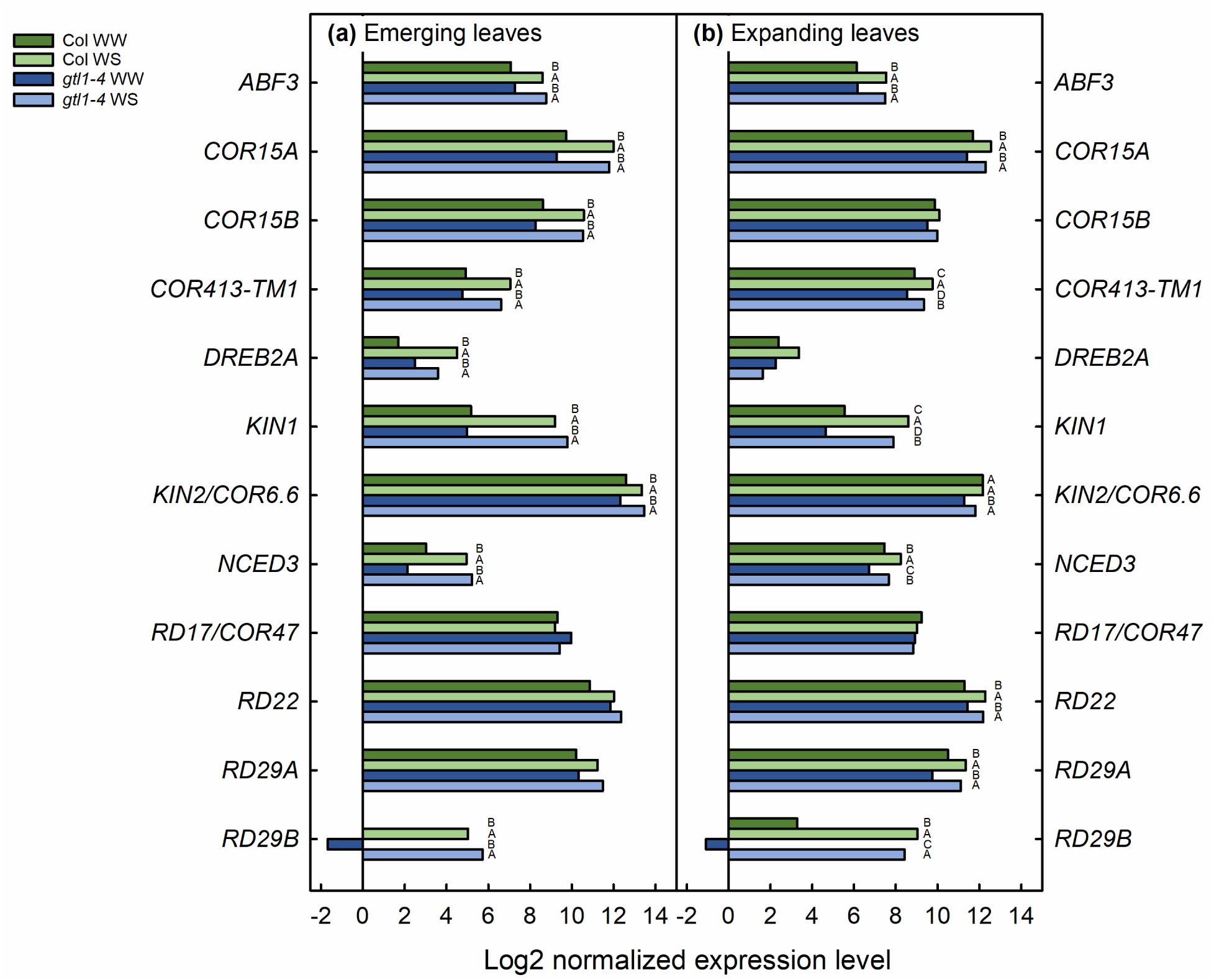
Expression of drought stress response markers in emerging (a) and expanding leaves (b). Data are shown as the log2-transformed normalized expression levels for three biological replicates per genotype in well-watered (WW) and water-stressed (WS) leaves. Statistically significant differences between genotype-treatment combinations are as given for *q*< 0.05 using the EBSeq-HMM package.

**Supplemental Figure 4.**
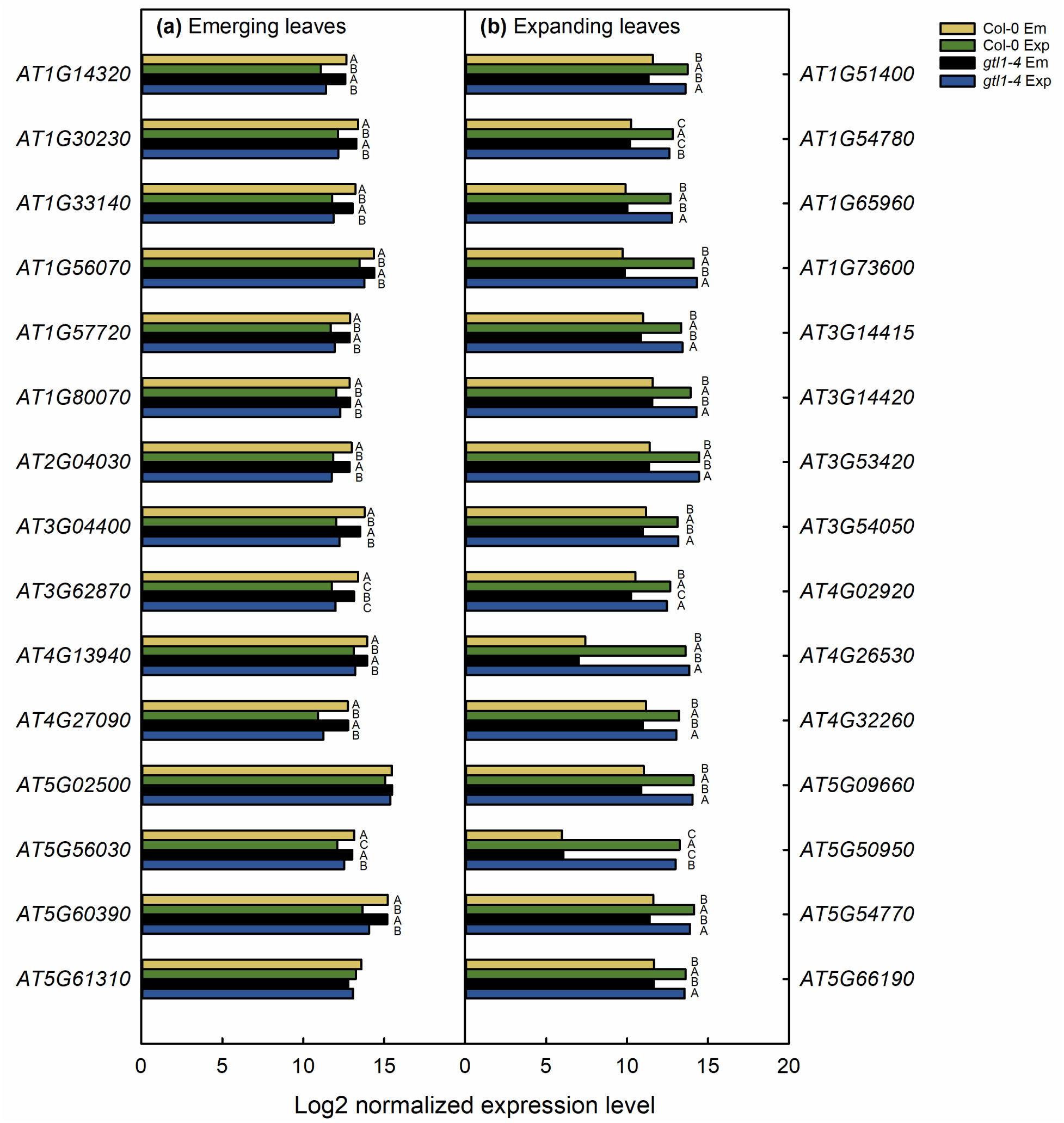
Expression of genes identified by Baerenfaller et al. (2012) as representative of emerging (a) and expanding (b) leaf stages. Data are shown as the log2-transformed normalized expression levels for three biological replicates per genotype-stage. Statistically significant differences between genotype-stage combinations are as given for *q*< 0.05 using the EBSeq-HMM package.

**Supplemental Figure 5.**
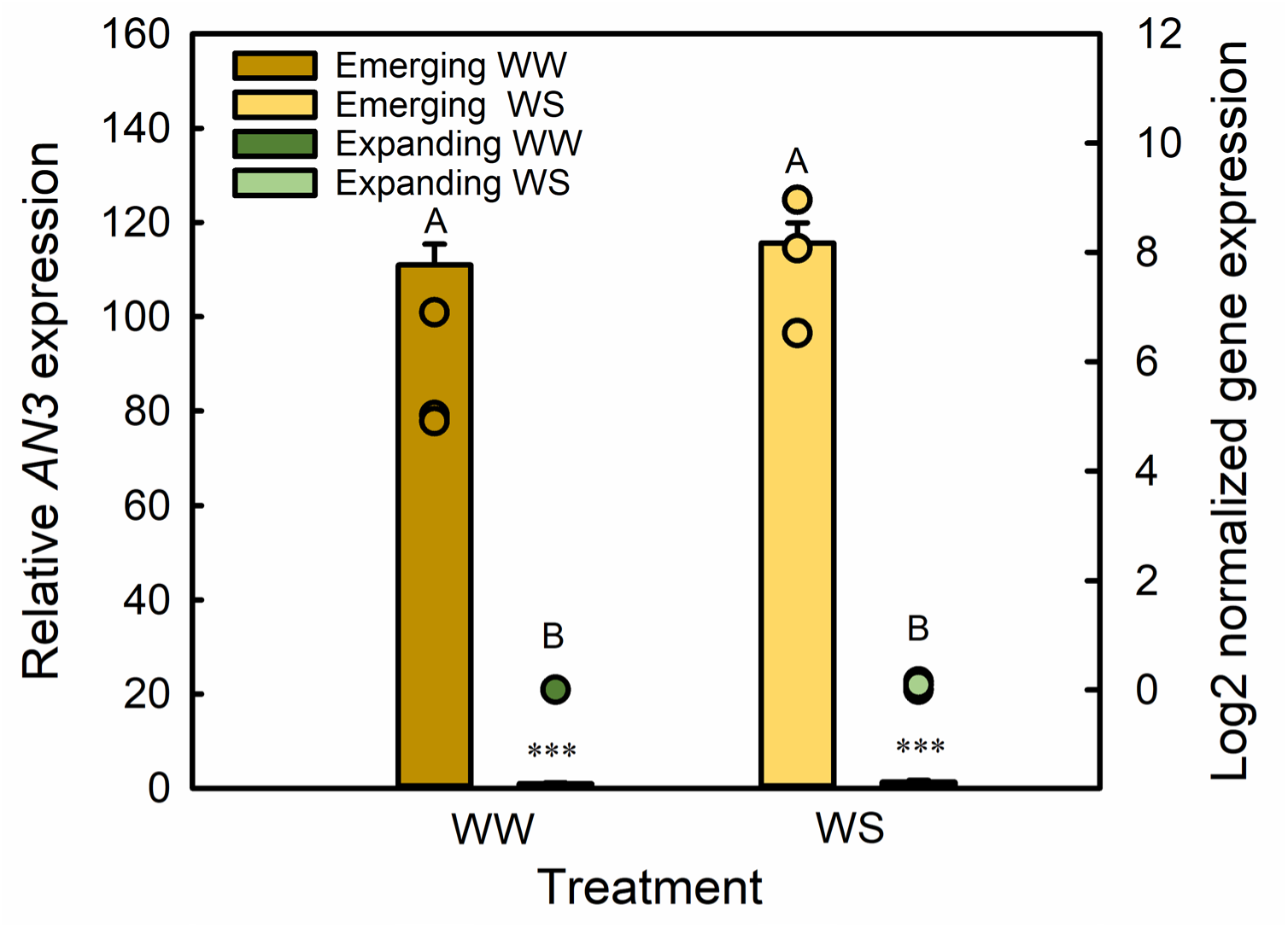
Expression of cell proliferation stage marker *AN3* as quantified by qPCR (bars ± standard error, left axis) or RNA-Seq (dots, right axis) in well-watered (WW) or water-stressed (WS) Col-0 leaves. Different letters above bars indicate mean separation as tested with EBSeq-HMM. qPCR data was compared against well-watered Col-0, and *** indicates a statistically significant difference at *P*< 0.001. *n=* 3.

**Supplemental Figure 6.**
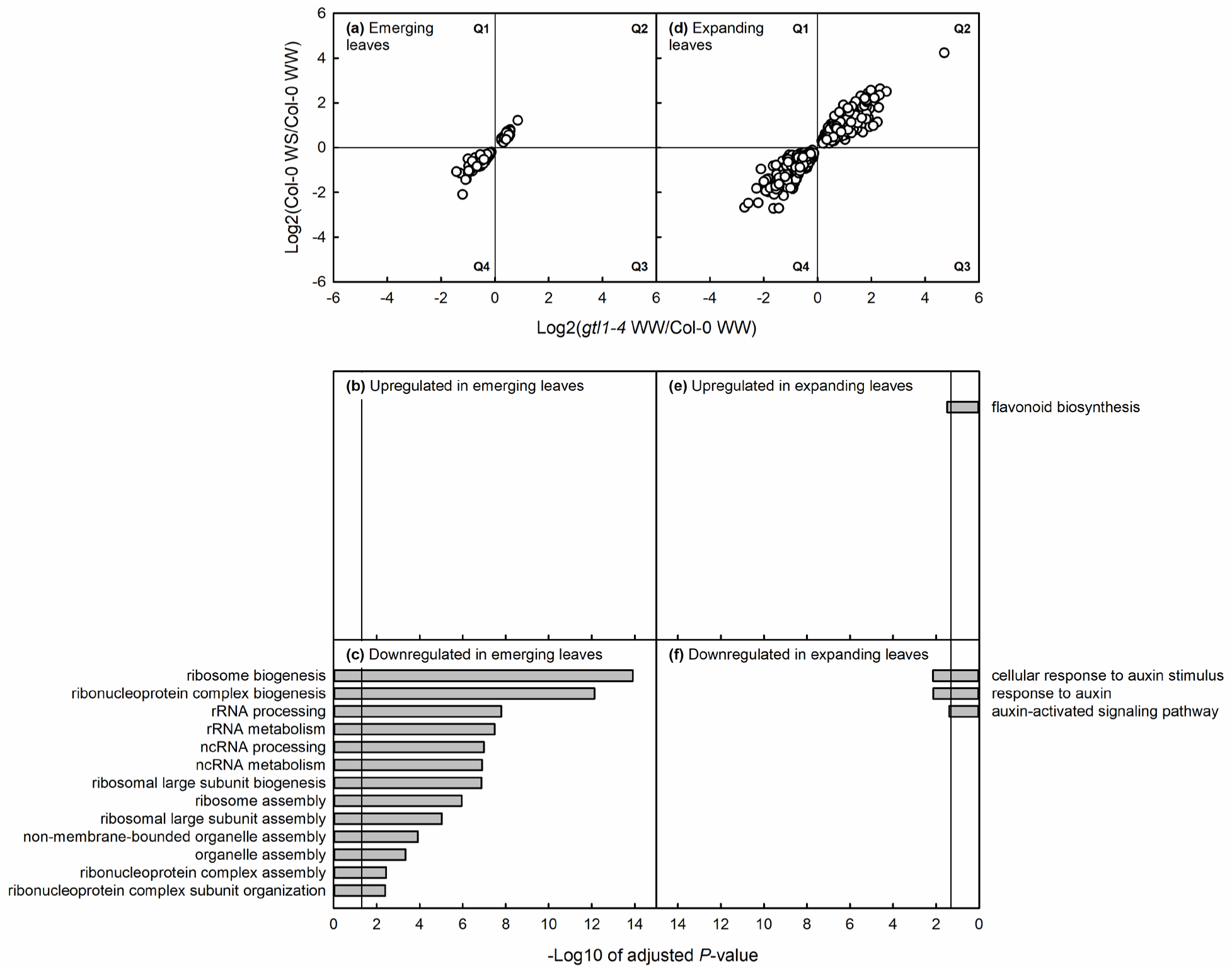
The log2-fold change of gene expression in well-watered (WW) *gtl1-4* relative to WW Col-0 against the FC in water-stressed (WS) Col-0 relative to WW Col-0 and the results of gene ontology overrepresentation analysis in emerging (a–c) and expanding (d–f) leaves. Data are plotted and shown for the 116 and 797 GTL1-primary-regulated genes (Figure 3), representing genes transcriptionally repressed or promoted by GTL1 or water deficit (Q2 and Q4, respectively). In GO analysis plots, all gene categories for which overrepresentation was statistically significant are shown. The vertical line in each plot represents the threshold of the false-discovery rate statistical significance.

**Supplemental Figure 7.**
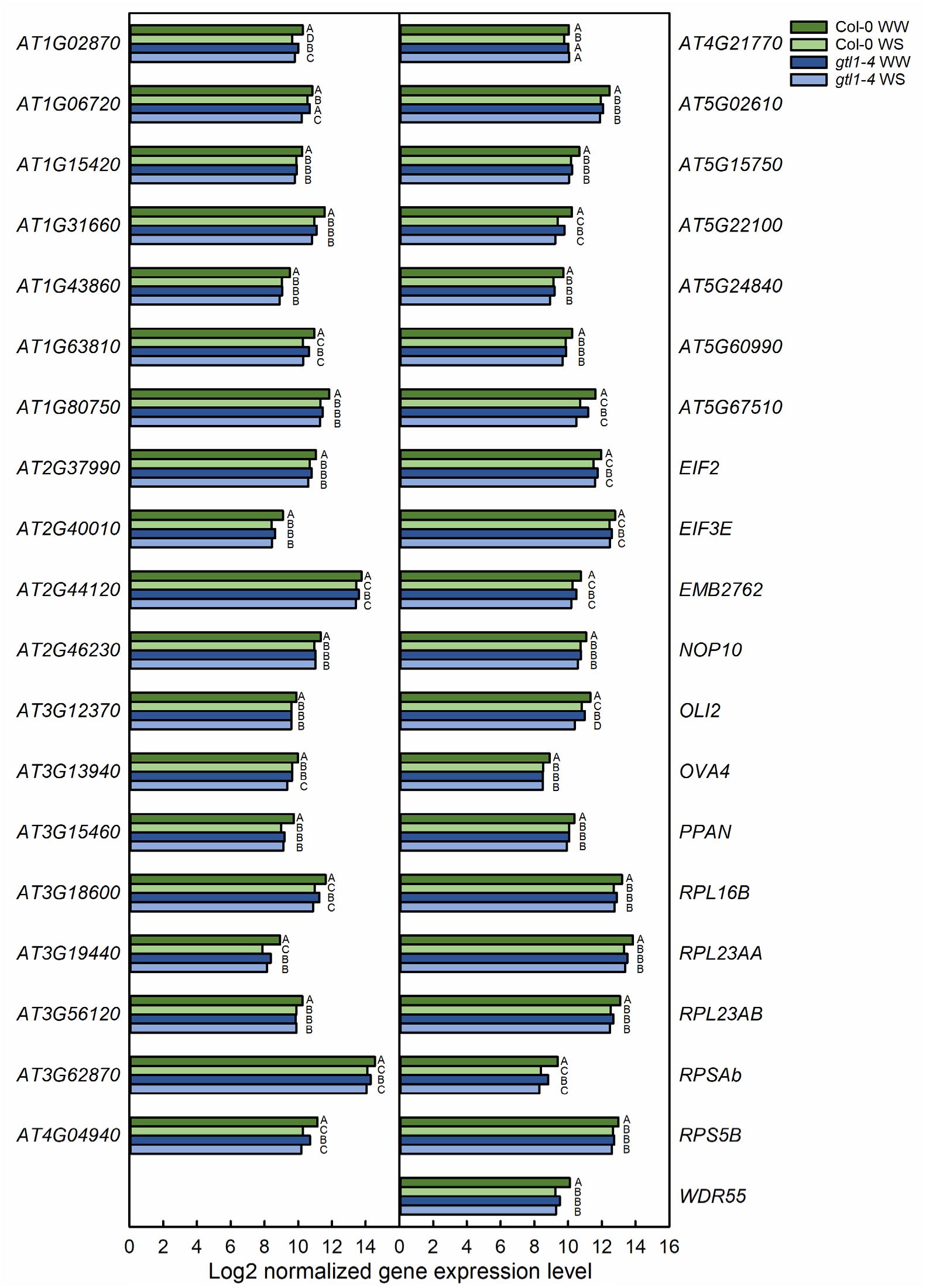
Expression of ribosome biogenesis genes as identified by gene ontology analysis (Fig. 4). Data are shown as the log2-transformed normalized expression levels for three biological replicates per genotype in well-watered (WW) and water-stressed (WS) leaves. Statistically significant differences between genotype-treatment combinations are as given for *q*< 0.05 using the EBSeq-HMM package.

## References

Alcázar, R., Bitrián, M., Bartels, D., Koncz, C., Altabella, T., & Tiburcio, A. F. (2011). Polyamine metabolic canalization in response to drought stress in *Arabidopsis* and the resurrection plant *Craterostigma plantagineum*. Plant Signaling and Behavior, 6, 243–250.

Alcázar, R., Cuevas, J. C., Patron, M., Altabella, T., & Tiburcio, A. F. (2006). Abscisic acid modulates polyamine metabolism under water stress in *Arabidopsis thaliana*. Physiologia Plantarum, 128, 448–455.

Alcázar, R., Planas, J., Saxena, T., Zarza, X., Bortolotti, C., Cuevas, J., Bitrián, M., Tiburcio, A. F., & Altabella, T. (2010). Putrescine accumulation confers drought tolerance in transgenic *Arabidopsis* plants over-expressing the homologous *Arginine decarboxylase 2* gene. Plant Physiology and Biochemistry, 48, 547–552.

Andriankaja, M., Dhondt, S., DeBodt, S., Vanhaeren, H., Coppens, F., DeMilde, L., Mühlenbock, P., Skirycz, A., Gonzalez, N., Beemster, G.T.S., & Inzé, D. (2012). Exit from proliferation during leaf development in *Arabidopsis thaliana*: A not-so-gradual process. Developmental Cell, 22, 64–78.

Anwar, R., Fatima, S., Mattoo, A. K., & Handa, A. K. (2019). Fruit architecture in polyamine-rich tomato germplasm is determined via a medley of cell cycle, cell expansion, and fruit shape genes. Plants, 8, 387.

Baerenfaller, K., Massonnet, C., Walsh, S., Baginsky, S., Buhlmann, P., Hennig, L., Hirsch-Hoffmann, M., Howell, K. A., Kahlau, S., Radziejwoski, A., Russenberger, D., Rutishauser, D., Small, I., Stekhoven, D., Sulpice, R., Svozil, J., Wuyts, N., Stitt, M., Hilson, P., Granier, C., & Gruissem, W. (2012). Systems-based analysis of Arabidopsis leaf growth reveals adaptation to water deficit. Molecular Systems Biology, 8, 606.

Baker, S. S., Wilhelm, K. S., & Thomashow, M. F. (1994). The 5’-region of *Arabidopsis thaliana cor15a* has *cis*-acting elements that confer cold-, drought- and ABA-regulated gene expression. Plant Molecular Biology, 24, 701–713.

Breuer, C., Kawamura, A., Ichikawa, T., Tominaga-Wada, R., Wada, T., Kondou, Y., Muto, S., Matsui, M., & Sugimoto, K. (2009). The trihelix transcription factor GTL1 regulates ploidy-dependent cell growth in the *Arabidopsis* trichome. The Plant Cell, 21, 2307–2322.

Breuer, C., Morohashi, K., Kawamura, A., Takahashi, N., Ishida, T., Umeda, M., Grotewold, E., & Sugimoto, K. (2012). Transcriptional repression of the APC/C activator CCS52A1 promotes active termination of cell growth. EMBO Journal, 31, 4488–4501.

Burow, M., & Halkier, B. A. (2017). How does a plant orchestrate defense in time and space? Using glucosinolates in Arabidopsis as case study. Current Opinion in Plant Biology, 38, 142– 147.

Chen, H., Li, Z., & Xiong, L. (2012). A plant microRNA regulates the adaptation of roots to drought stress. FEBS Letters, 586, 1742–1747.

Chen, J., Nolan, T. M., Ye, H., Zhang, M., Tong, H., Xin, P., Chu, J., Chu, C., Li, Z., & Yina, Y. (2017). Arabidopsis WRKY46, WRKY54, and WRKY70 transcription factors are involved in brassinosteroid-regulated plant growth and drought responses. The Plant Cell, 29, 1425–1439.

Chen, Y., Dubois, M., Vermeersch, M., Inzé, D., & Vanhaeren, H. (2021). Distinct cellular strategies determine sensitivity to mild drought of Arabidopsis natural accessions. Plant Physiology, 186, 1171–1185.

D’Alessandro, S., Golin, S., Hardtke, C. S., Lo Schiavo, F., & Zottini, M. (2015). The co-chaperone p23 controls root development through the modulation of auxin distribution in the *Arabidopsis* root meristem. Journal of Experimental Botany, 66, 5113–5122.

Ding, X., Wang, X., Li, Q., Yu, L., Song, Q., Gai, J., & Yang, S. (2019). Metabolomics studies on cytoplasmic male sterility during flower bud development in soybean. International Journal of Molecular Sciences, 20, 2869.

Donahue, J. L., Alford, S. R., Torabinejad, J., Kerwin, R. E., Nourbakhsh, A., Keith Ray, W., Hernick, M., Huang, X., Lyons, B. M., Hein, P. P., & Gillaspy, G. E. (2010). The *Arabidopsis thaliana myo*-inositol 1-phosphate synthase1 gene is required for *myo*-inositol synthesis and suppression of cell death. The Plant Cell, 22, 888–903.

Donnelly, P. M., Bonetta, D., Tsukaya, H., Dengler, R. E., & Dengler, N. G. (1999). Cell cycling and cell enlargement in developing leaves of *Arabidopsis*. Developmental Biology, 215, 407– 419.

Endo, A., Sawada, Y., Takahashi, H., Okamoto, M., Ikegami, K., Koiwai, H., Seo, M., Toyomasu, T., Mitsuhashi, W., Shinozaki, K., Nakazono, M., Kamiya, Y., Koshiba, T., & Nambara, E. (2008). Drought induction of Arabidopsis 9-cis-epoxycarotenoid dioxygenase occurs in vascular parenchyma cells. Plant Physiology, 147, 1984–1993.

Feng, Z. J., Xu, S. C., Liu, N., Zhang, G. W., Hu, Q. Z., Xu, Z. S., & Gong, Y. M. (2018). Identification of the AQP members involved in abiotic stress responses from *Arabidopsis*. Gene, 646, 64–73.

Figueroa, N., Lodeyro, A. F., Carrillo, N., & Gómez, R. (2021). Meta-analysis reveals key features of the improved drought tolerance of plants overexpressing NAC transcription factors. Environmental and Experimental Botany, 186, 104449.

Franceschetti, M., Hanfrey, C., Scaramagli, S., Torrigiani, P., Bagni, N., Burtin, D., & Michael, A. J. (2001). Characterization of monocot and dicot plant S-adenosyl-l-methionine decarboxylase gene families including identification in the mRNA of a highly conserved pair of upstream overlapping open reading frames. Biochemical Journal, 353, 403–409.

Fujikura, U., Horiguchi, G., Ponce, M. R., Micol, J. L., & Tsukaya, H. (2009). Coordination of cell proliferation and cell expansion mediated by ribosome-related processes in the leaves of *Arabidopsis thaliana*. The Plant Journal, 59, 499–508.

Geisler, M., Nadeau, J., & Sack, F. D. (2000). Oriented asymmetric divisions that generate the stomatal spacing pattern in Arabidopsis are disrupted by the *too many mouths* mutation. The Plant Cell, 12, 2075–2086.

Georgii, E., Jin, M., Zhao, J., Kanawati, B., Schmitt-Kopplin, P., Albert, A., Winkler, J. B., & Schäffner, A. R. (2017). Relationships between drought, heat and air humidity responses revealed by transcriptome-metabolome co-analysis. BMC Plant Biology, 17, 120.

Gonzalez, N., Vanhaeren, H., & Inze, D. (2012). Leaf size control: Complex coordination of cell division and expansion. Trends in Plant Science, 17, 1360–1385.

Hanfrey, C., Sommer, S., Mayer, M. J., Burtin, D., & Michael, A. J. (2001). *Arabidopsis* polyamine biosynthesis: Absence of ornithine decarboxylase and the mechanism of arginine decarboxylase activity. The Plant Journal, 27, 551–560.

Harb, A., Krishnan, A., Ambavaram, M. M. R., & Pereira, A. (2010). Molecular and physiological analysis of drought stress in Arabidopsis reveals early responses leading to acclimation in plant growth. Plant Physiology, 154, 1254–1271.

Havaux, M., & Kloppstech, K. (2001). The protective functions of carotenoid and flavonoid pigments against excess visible radiation at chilling temperature investigated in *Arabidopsis npq* and *tt* mutants. Planta, 213, 953–966.

Herrero, A., Sanllorente, S., Reguera, C., Ortiz, M. C., & Sarabia, L. A. (2016). A new multiresponse optimization approach in combination with a D-Optimal experimental design for the determination of biogenic amines in fish by HPLC-FLD. Analytica Chimica Acta, 945, 31– 38.

Himmelbach, A., Hoffmann, T., Leube, M., Höhener, B., & Grill, E. (2002). Homeodomain protein ATHB6 is a target of the protein phosphatase ABI1 and regulates hormone responses in *Arabidopsis*. The EMBO Journal, 21, 3029–3038.

Horiguchi, G., Kim, G. T., & Tsukaya, H. (2005). The transcription factor AtGRF5 and the transcription coactivator AN3 regulate cell proliferation in leaf primordia of *Arabidopsis thaliana*. The Plant Journal, 43, 68–78.

Ichino, T., Fuji, K., Ueda, H., Takahashi, H., Koumoto, Y., Takagi, J., Tamura, K., Sasaki, R., Aoki, K., Shimada, T., & Hara-Nishimura, I. (2014). GFS9/TT9 contributes to intracellular membrane trafficking and flavonoid accumulation in *Arabidopsis thaliana*. The Plant Journal, 80, 410–423.

Ivanov, I. P., Atkins, J. F., & Michael, A. J. (2010). A profusion of upstream open reading frame mechanisms in polyamine-responsive translational regulation. Nucleic Acids Research, 38, 353– 359.

Kang, J., & Dengler, N. (2004). Vein pettern development n adult leaves of *Arabidopsis thaliana*. International Journal of Plant Science, 165, 231–242.

Khan, M., Imran, Q. M., Shahid, M., Mun, B. G., Lee, S. U., Khan, M. A., Hussain, A., Lee, I. J., & Yun, B. W. (2019). Nitric oxide-induced *AtAO3* differentially regulates plant defense and drought tolerance in *Arabidopsis thaliana*. BMC Plant Biology, 19, 602.

Kodaira, K. S., Qin, F., Tran, L. S. P., Maruyama, K., Kidokoro, S., Fujita, Y., Shinozaki, K., & Yamaguchi-Shinozaki, K. (2011). Arabidopsis Cys2/His2 zinc-finger proteins AZF1 and AZF2 negatively regulate abscisic acid-repressive and auxin-inducible genes under abiotic stress conditions. Plant Physiology, 157, 742–756.

Koiwai, H., Nakaminami, K., Seo, M., Mitsuhashi, W., Toyomasu, T., & Koshiba, T. (2004). Tissue-specific localization of an abscisic acid biosynthetic enzyme, AAO3, in Arabidopsis. Plant Physiology, 134, 1697–1707.

Komili, S., Farny, N. G., Roth, F. P., & Silver, P. A. (2007). Functional specificity among ribosomal proteins regulates gene expression. Cell, 131, 557–571.

Kumar, M. N., Hsieh, Y. F., & Verslues, P. E. (2015). At14a-Like1 participates in membrane-associated mechanisms promoting growth during drought in *Arabidopsis thaliana*. Proceedings of the National Academy of Sciences of the United States of America, 112, 10545–10550.

Kumari, A., Jewaria, P. K., Bergmann, D. C., & Kakimoto, T. (2014). Arabidopsis reduces growth under osmotic stress by decreasing SPEECHLESS protein. Plant & Cell Physiology, 55, 2037–2046.

Larkin, J. C., Young, N., Prigge, M., & Marks, M. D. (1996). The control of trichome spacing and number in *Arabidopsis*. Development, 122, 997–1005.

Lechner, E., Leonhardt, N., Eisler, H., Parmentier, Y., Alioua, M., Jacquet, H., Leung, J., & Genschik, P. (2011). MATH/BTB CRL3 receptors target the homeodomain-leucine zipper ATHB6 to modulate abscisic acid signaling. Developmental Cell, 21, 1116–1128.

Leng, N., Li, Y., McIntosh, B. E., Nguyen, B. K., Duffin, B., Tian, S., Thomson, J. A., Dewey, C. N., Stewart, R., & Kendziorski, C. (2015). EBSeq-HMM: A Bayesian approach for identifying gene-expression changes in ordered RNA-seq experiments. Bioinformatics, 31, 2614–2622.

Li, J., Zhou, X., Xiong, C., Zhou, H., Li, H., & Ruan, C. (2023). Yellowhorn Xso-miR5149-*XsGTL1* enhances water-use efficiency and drought tolerance by regulating leaf morphology and stomatal density. International Journal of Biological Macromolecules, 237, 124060.

Li, T., Zhang, Y., Liu, Y., Li, X., Hao, G., Han, Q., Dirk, L. M. A., Downie, A. B., Ruan, Y. L., Wang, J., Wang, G., & Zhao, T. (2020). Raffinose synthase enhances drought tolerance through raffinose synthesis or galactinol hydrolysis in maize and *Arabidopsis* plants. Journal of Biological Chemistry, 295, 8064–8077.

Li, W. X., Oono, Y., Zhu, J., He, X. J., Wu, J. M., Iida, K., Lu, X. Y., Cui, X., Jin, H., & Zhu, J. K. (2008). The *Arabidopsis* NFYA5 transcription factor is regulated transcriptionally and posttranscriptionally to promote drought resistance. The Plant Cell, 20, 2238–2251.

Liu, Q., Kasuga, M., Sakuma, Y., Abe, H., Miura, S., Yamaguchi-Shinozaki, K., & Shinozaki, K. (1998). Two transcription factors, DREB1 and DREB2, with an EREBP/AP2 DNA binding domain separate two cellular signal transduction pathways in drought- and low-temperature-responsive gene expression, respectively, in Arabidopsis. The Plant Cell, 10, 1391–1406.

Liu, Q., Wang, Z., Yu, S., Li, W., Zhang, M., Yang, J., Li, D., Yang, J., & Li, C. (2021). PumiR172d regulates stomatal density and water-use efficiency via targeting *PuGTL1* in poplar. Journal of Experimental Botany, 72, 1370–1383.

Manna, M., Thakur, T., Chirom, O., Mandlik, R., Deshmukh, R., & Salvi, P. (2021). Transcription factors as key molecular target to strengthen the drought stress tolerance in plants. Physiologia Plantarum, 172, 847–868.

Martinez-Seidel, F., Beine-Golovchuk, O., Hsieh, Y. C., & Kopka, J. (2020). Systematic review of plant ribosome heterogeneity and specialization. Frontiers in Plant Science, 11, 948.

Merret, R., Carpentier, M. C., Favory, J. J., Picart, C., Descombin, J., Bousquet-Antonelli, C., Tillard, P., Lejay, L., Deragon, J. M., & Charng, Y. Y. (2017). Heat shock protein HSP101 affects the release of ribosomal protein mRNAs for recovery after heat shock. Plant Physiology, 174, 1216–1225.

Miller, G., Schlauch, K., Tam, R., Cortes, D., Torres, M. A., Shulaev, V., Dangl, J. L., & Mittler, R. (2009). The plant NADPH oxidase RBOHD mediates rapid systemic signaling in response to diverse stimuli. Science Signaling, 2, ra45.

Mudgil, Y., Uhrig, J. F., Zhou, J., Temple, B., Jiang, K., & Jones, A. M. (2009). *Arabidopsis* N-MYC DOWNREGULATED-LIKE1, a positive regulator of auxin transport in a G protein-mediated pathway. The Plant Cell, 21, 3591–3609.

Nagata, T., Todoriki, S., Masumizu, T., Suda, I., Furuta, S., Du, Z., & Kikuchi, S. (2003). Levels of active oxygen species are controlled by ascorbic acid and anthocyanin in *Arabidopsis*. Journal of Agricultural and Food Chemistry, 51, 2992–2999.

Nakabayashi, R., Yonekura-Sakakibara, K., Urano, K., Suzuki, M., Yamada, Y., Nishizawa, T., Matsuda, F., Kojima, M., Sakakibara, H., Shinozaki, K., Michael, A. J., Tohge, T., Yamazaki, M., & Saito, K. (2014). Enhancement of oxidative and drought tolerance in *Arabidopsis* by overaccumulation of antioxidant flavonoids. The Plant Journal, 77, 367–379.

Nakano, R. T., Piślewska-Bednarek, M., Yamada, K., Edger, P. P., Miyahara, M., Kondo, M., Böttcher, C., Mori, M., Nishimura, M., Schulze-Lefert, P., Hara-Nishimura, I., & Bednarek, P. (2017). PYK10 myrosinase reveals a functional coordination between endoplasmic reticulum bodies and glucosinolates in *Arabidopsis thaliana*. The Plant Journal, 89, 204–220.

Omidbakhshfard, M. A., Sokolowska, E. M., Di Vittori, V., Perez de Souza, L., Kuhalskaya, A., Brotman, Y., Alseekh, S., Fernie, A. R., & Skirycz, A. (2021). Multi-omics analysis of early leaf development in *Arabidopsis thaliana*. Patterns, 2, 100235.

Pauwels, L., Morreel, K., De Witte, E., Lammertyn, F., Van Montagu, M., Boerjan, W., Inzé, D., & Goossens, A. (2008). Mapping methyl jasmonate-mediated transcriptional reprogramming of metabolism and cell cycle progression in cultured *Arabidopsis* cells. Proceedings of the National Academy of Sciences of the United States of America, 105, 1380–1385.

Pfaffl, M. W. (2001). A new mathematical model for relative quantification in real-time RT-PCR. Nucleic Acids Research, 29, e45.

Quirino, B. F., Reiter, W. -D., & Amasino, R. D. (2001). One of two tandem *Arabidopsis* genes homologous to monosaccharide transporters is senescence-associated. Plant Molecular Biology, 46, 447–457.

Rabanal, F. A., Mandáková, T., Soto-Jiménez, L. M., Greenhalgh, R., Parrott, D. L., Lutzmayer, S., Steffen, J. G., Nizhynska, V., Mott, R., Lysak, M. A., Clark, R. M., & Nordborg, M. (2017). Epistatic and allelic interactions control expression of ribosomal RNA gene clusters in *Arabidopsis thaliana*. Genome Biology, 18, 75.

Rosado, A., Sohn, E. J., Drakakaki, G., Pan, S., Swidergal, A., Xiong, Y., Kang, B. H., Bressan, R. A., & Raikhel, N. V. (2010). Auxin-mediated ribosomal biogenesis regulates vacuolar trafficking in *Arabidopsis*. The Plant Cell, 22, 143–158.

Roumani, M., Besseau, S., Gagneul, D., Robin, C., & Larbat, R. (2021). Phenolamides in plants: An update on their function, regulation, and origin of their biosynthetic enzymes. Journal of Experimental Botany, 72, 2334–2355.

Rymaszewski, W., Vile, D., Bediee, A., Dauzat, M., Luchaire, N., Kamrowska, D., Granier, C., & Hennig, J. (2017). Stress-related gene expression reflects morphophysiological responses to water deficit. Plant Physiology, 174, 1913–1930.

Seo, M., Aoki, H., Koiwai, H., Kamiya, Y., Nambara, E., & Koshiba, T. (2004). Comparative studies on the *Arabidopsis* aldehyde oxidase (*AAO*) gene family revealed a major role of *AAO3* in ABA biosynthesis in seeds. Plant and Cell Physiology, 45, 1694–1703.

Seo, M., Peeters, A. J. M., Koiwai, H., Oritani, T., Marion-Poll, A., Zeevaart, J. A. D., Koornneef, M., Kamiya, Y., & Koshiba, T. (2000). The *Arabidopsis* aldehyde oxidase 3 (*AAO3*) gene product catalyzes the final step in abscisic acid biosynthesis in leaves. Proceedings of the National Academy of Sciences of the United States of America, 97, 12908–12913.

Shi, H., Chen, L., Ye, T., Liu, X., Ding, K., & Chan, Z. (2014). Modulation of auxin content in Arabidopsis confers improved drought stress resistance. Plant Physiology and Biochemistry, 82, 209–217.

Shibata, M., Breuer, C., Kawamura, A., Clark, N. M., Rymen, B., Braidwood, L., Morohashi, K., Busch, W., Benfey, P. N., Sozzani, R., & Sugimoto, K. (2018). GTL1 and DF1 regulate root hair growth through transcriptional repression of *ROOT HAIR DEFECTIVE 6-LIKE 4* in *Arabidopsis*. Development, 145, 159707.

Shin, D. J., Min, J. H., Van Nguyen, T., Kim, Y. M., & Kim, C. S. (2019). Loss of *Arabidopsis Halotolerance 2-like* (*AHL*), a 3′-phosphoadenosine-5′-phosphate phosphatase, suppresses insensitive response of *Arabidopsis thaliana ring zinc finger 1* (*atrzf1*) mutant to abiotic stress. Plant Molecular Biology, 99, 363–377.

Skirycz, A., De Bodt, S., Obata, T., De Clercq, I., Claeys, H., De Rycke, R., Andriankaja, M., Van Aken, O., Van Breusegem, F., Fernie, A. R., & Inzé, D. (2010) Developmental stage specificity and the role of mitochondrial metabolism in the response of Arabidopsis leaves to prolonged mild osmotic stress. Plant Physiology, 152, 226–244.

Skirycz, A., Claeys, H., De Bodt, S., Oikawa, A., Shinoda, S., Andriankaja, M., Maleux, K., Eloy, N. B., Coppens, F., Yoo, S. -D., Saito, K., & Inzé, D. (2011). Pause-and-stop: The effects of osmotic stress on cell proliferation during early leaf development in *Arabidopsis* and a role for ethylene signaling in cell cycle arrest. The Plant Cell, 23, 1876–1888.

Slawinski, L., Israel, A., Artault, C., Thibault, F., Atanassova, R., Laloi, M., & Dédaldéchamp, F. (2021). Responsiveness of early response to dehydration six-like transporter genes to water deficit in *Arabidopsis thaliana* leaves. Frontiers in Plant Science, 12, 708876.

Sormani, R., Masclaux-Daubresse, C., Daniele-Vedele, F., & Chardon, F. (2011). Transcriptional regulation of ribosome components are determined by stress according to cellular compartments in *Arabidopsis thaliana*. PLoS ONE, 6, e28070.

Sperdouli, I., Moustaka, J., Ouzounidou, G., & Moustakas, M. (2021). Leaf age-dependent photosystem II photochemistry and oxidative stress responses to drought stress in *Arabidopsis thaliana* are modulated by flavonoid accumulation. Molecules, 26, 4157.

Sperdouli, I. & Moustakas, M. (2014). Leaf developmental stage modulates metabolite accumulation and photosynthesis contributing to acclimation of *Arabidopsis thaliana* to water deficit. Journal of Plant Research, 127, 481–489.

Spribille, R. & Forkmann, G. (1982). Chalcone synthesis and hydroxylation of flavonoids in 3’-position with enzyme preparations from flowers of *Dianthus caryophyllus* L. (carnation). Planta, 155, 176–182.

Staswick, P. E., Serban, B., Rowe, M., Tiryaki, I., Maldonado, M. T., Maldonado, M. C., & Suza, W. (2005). Characterization of an Arabidopsis enzyme family that conjugates amino acids to indole-3-acetic acid. The Plant Cell, 17, 616–627.

Sun, Y., Li, H., & Huang, J. R. (2012). *Arabidopsis* TT19 functions as a carrier to transport anthocyanin from the cytosol to tonoplasts. Molecular Plant, 5, 387–400.

Takahashi, Y., Cong, R., Sagor, G. H. M., Niitsu, M., Berberich, T., & Kusano, T. (2010). Characterization of five polyamine oxidase isoforms in *Arabidopsis thaliana*. Plant Cell Reports, 29, 955–965.

Tao, L. Z., Cheung, A. Y., Nibau, C., & Wu, H. M. (2005). RAC GTPases in tobacco and Arabidopsis mediate auxin-induced formation of proteolytically active nuclear protein bodies that contain AUX/IAA proteins. The Plant Cell, 17, 2369–2383.

Tohge, T., Matsui, K., Ohme-Takagi, M., Yamazaki, M., & Saito, K. (2005). Enhanced radical scavenging activity of genetically modified *Arabidopsis* seeds. Biotechnology Letters, 27, 297– 303.

Tsuge, T., Tsukaya, H., & Uchimiya, H. (1996). Two independent and polarized processes of cell elongation regulate leaf blade expansion in *Arabidopsis thaliana* (L.) Heynh. Development, 122, 1589–1600.

Urano, K., Yoshiba, Y., Nanjo, T., Ito, T., Yamaguchi-Shinozaki, K., & Shinozaki, K. (2004). *Arabidopsis* stress-inducible gene for arginine decarboxylase *AtADC2* is required for accumulation of putrescine in salt tolerance. Biochemical and Biophysical Research Communications, 313, 369–375.

Völz, R., Kim, S. K., Mi, J., Mariappan, K. G., Guo, X., Bigeard, J., Alejandro, S., Pflieger, D., Rayapuram, N., Al-Babili, S., & Hirt, H. (2018). The trihelix transcription factor GT2-like 1 (GTL1) promotes salicylic acid metabolism, and regulates bacterial-triggered immunity. PLoS Genetics, 14, e1007708.

Wade, H. K., Sohal, A. K., & Jenkins, G. I. (2003). Arabidopsis *ICX1* is a negative regulator of several pathways regulating flavonoid biosynthesis genes. Plant Physiology, 131, 707–715.

Wang, C., Li, J., Ma, M., Lin, Z., Hu, W., Lin, W., & Zhang, P. (2021). Structural and biochemical insights into two BAHD acyltransferases (*At*SHT and *At*SDT) involved in phenolamide biosynthesis. Frontiers in Plant Science, 11, 610118.

Wang, F., Kong, W., Wong, G., Fu, L., Peng, R., Li, Z., & Yao, Q. (2016). *AtMYB12* regulates flavonoids accumulation and abiotic stress tolerance in transgenic *Arabidopsis thaliana*. Molecular Genetics and Genomics, 291, 1545–1559.

Wang, H., Zhou, Y., Gilmer, S., Whitwill, S., & Fowke, L. C. (2000) Expression of the plant cyclin-dependent kinase inhibitor ICK1 affects cell division, plant growth and morphology. The Plant Journal, 24, 613–623.

Wang, H., Fowke, L. C., & Crosby, W.L. (1997) A plant cyclin-dependent kinase inhibitor gene. Nature, 386, 451–452.

Watson, M. B., Emory, K. K., Piatak, R. M., & Malmberg, R. L. (1998). Arginine decarboxylase (polyamine synthesis) mutants of *Arabidopsis thaliana* exhibit altered root growth. The Plant Journal, 13, 231–239.

Weng, H., Yoo, C. Y., Gosney, M. J., Hasegawa, P. M., & Mickelbart, M. V. (2012). Poplar GTL1 is a Ca^2+^/calmodulin-binding transcription factor that functions in plant water use efficiency and drought tolerance. PLoS ONE, 7, e32925.

Wieckowski, Y. & Schiefelbein, J. (2012). Nuclear ribosome biogenesis mediated by the DIM1A rRNA dimethylase is required for organized root growth and epidermal patterning in *Arabidopsis*. The Plant Cell, 24, 2839–2856.

Wu, H., Fu, B., Sun, P., Xiao, C., & Liu, J. H. (2016). A NAC transcription factor represses putrescine biosynthesis and affects drought tolerance. Plant Physiology, 172, 1532–1547.

Xie, Z., Li, D., Wang, L., Sack, F. D., & Grotewold, E. (2010). Role of the stomatal development regulators FLP/MYB88 in abiotic stress responses. The Plant Journal, 64, 731–739.

Xie, Z., Nolan, T. M., Jiang, H., & Yin, Y. (2019). AP2/ERF transcription factor regulatory networks in hormone and abiotic stress responses in *Arabidopsis*. Frontiers in Plant Science, 10, 228.

Xu, W., Grain, D., Bobet, S., Le Gourrierec, J., Thévenin, J., Kelemen, Z., Lepiniec, L., & Dubos, C. (2014). Complexity and robustness of the flavonoid transcriptional regulatory network revealed by comprehensive analyses of MYB-bHLH-WDR complexes and their targets in Arabidopsis seed. New Phytologist, 202, 132–144.

Xu, Z., Mahmood, K., & Rothstein, S. J. (2017). ROS induces anthocyanin production via late biosynthetic genes and anthocyanin deficiency confers the hypersensitivity to ROS-generating stresses in *Arabidopsis*. Plant and Cell Physiology, 58, 1364–1377.

Yamada, K., Nagano, A. J., Nishina, M., Hara-Nishimura, I., & Nishimura, M. (2008). NAI2 is an endoplasmic reticulum body component that enables ER body formation in *Arabidopsis thaliana*. The Plant Cell, 20, 2529–2540.

Yamaguchi-Shinozaki, K. & Shinozaki, K. (1993). Characterization of the expression of a desiccation-responsive *rd29* gene of *Arabidopsis thaliana* and analysis of its promoter in transgenic plants. Molecular and General Genetics, 236, 331–340.

Yoo, C. Y., Mano, N., Finkler, A., Weng, H., Day, I. S., Reddy, A. S. N., Poovaiah, B. W., Fromm, H., Hasegawa, P. M., & Mickelbart, M. V. (2019). A Ca^2+^/CaM-regulated transcriptional switch modulates stomatal development in response to water deficit. Scientific Reports, 9, 12282.

Yoo, C. Y., Pence, H. E., Jin, J. B., Miura, K., Gosney, M. J., Hasegawa, P. M., & Mickelbart, M. V. (2010). The *Arabidopsis* GTL1 transcription factor regulates water use efficiency and drought tolerance by modulating stomatal density via transrepression of *SDD1*. The Plant Cell, 22, 4128–4141.

Yu, Z., Zhang, P., Lin, W., Zheng, X., Cai, M., & Peng, C. (2019). Sequencing of anthocyanin synthesis-related enzyme genes and screening of reference genes in leaves of four dominant subtropical forest tree species. Gene, 716, 144024.

Yuan, X., Xu, P., Yu, Y., & Xiong, Y. (2020). Glucose-TOR signaling regulates PIN2 stability to orchestrate auxin gradient and cell expansion in *Arabidopsis* root. Proceedings of the National Academy of Sciences of the United States of America, 117, 32223–32225.

Zhang, Q., Zhai, J., Chen, G., Lin, W., & Peng, C. (2019). The changing distribution of anthocyanin in *Mikania micrantha* leaves as an adaption to low-temperature environments. Plants, 8, 456.

Zhang, X. -H., Zheng, X. -T., Sun, B. -Y., Peng, C. -L., & Chow, W. S. (2018). Over-expression of the *CHS* gene enhances resistance of *Arabidopsis* leaves to high light. Environmental and Experimental Botany, 154, 33–43.

Zhao, P. X., Zhang, J., Chen, S. Y., Wu, J., Xia, J. Q., Sun, L. Q., Ma, S. S., & Xiang, C. B. (2021). Arabidopsis MADS-box factor AGL16 is a negative regulator of plant response to salt stress by downregulating salt-responsive genes. New Phytologist, 232, 2418–2439.

Zheng, X., Liu, H., Ji, H., Wang, Y., Dong, B., Qiao, Y., Liu, M., & Li, X. (2016). The wheat GT factor *TaGT2L1D* negatively regulates drought tolerance and plant development. Scientific Reports, 6, 27042.

Zhou, D. -X. (1999). Regulatory mechanism of plant gene transcription by GT-elements and GT-factors. Trends in Plant Science, 4, 210–214.

Zhou, L., Liu, Z., Liu, Y., Kong, D., Li, T., Yu, S., Mei, H., Xu, X., Liu, H., Chen, L., & Luo, L. (2016). A novel gene *OsAHL1* improves both drought avoidance and drought tolerance in rice. Scientific Reports, 6, 30264.

Zhou, X. F., Jin, Y. H., Yoo, C. Y., Lin, X. L., Kim, W. Y., Yun, D. J., Bressan, R. A., Hasegawa, P. M., & Jin, J. B. (2013). CYCLIN H;1 regulates drought stress responses and blue light-induced stomatal opening by inhibiting reactive oxygen species accumulation in Arabidopsis. Plant Physiology, 162, 1030–1041.

